# Structural landscape of AAA+ ATPase motor states in the substrate-degrading human 26S proteasome reveals conformation-specific binding of TXNL1

**DOI:** 10.1101/2024.11.08.622731

**Authors:** Connor Arkinson, Christine L. Gee, Zeyuan Zhang, Ken C. Dong, Andreas Martin

## Abstract

The 26S proteasome targets many cellular proteins for degradation during general homeostasis, protein quality control, and the regulation of vital processes. A broad range of proteasome-interacting cofactors thereby modulates these functions and aids in substrate degradation. Here, we solved several high-resolution structures of the redox active cofactor TXNL1 bound to the human 26S proteasome at saturating and sub-stoichiometric concentrations by time resolved cryo-EM. We identified distinct binding modes of TXNL1 that depend on the proteasome conformational and ATPase motor states. Together with biophysical and biochemical experiments, our structural studies reveal that the resting-state proteasome prior to substrate engagement with the ATPase motor binds TXNL1 with low affinity and in variable positions on top of the Rpn11 deubiquitinase. In contrast, the actively degrading proteasome shows additional interactions leading to high-affinity TXNL1 binding, whereby TXNL1’s C-terminal tail covers the catalytic groove of the Rpn11 deubiquitinase and coordinates the active-site Zn^2+^. Furthermore, these cryo-EM structures of the degrading proteasome capture the ATPase hexamer in all registers of spiral-staircase arrangements and thus visualize the complete ATP-hydrolysis cycle of the AAA+ motor, indicating temporally asymmetric hydrolysis and conformational changes in bursts during mechanical substrate unfolding and translocation. Remarkably, we catch the proteasome in the act of unfolding the beta-barrel mEos3.2 substrate while the ATPase hexamer is in a particular spiral staircase register. Our findings challenge current models for protein translocation through hexameric AAA+ motors and reveal how the proteasome uses its distinct but broad range of conformational states to coordinate cofactor binding and substrate processing.

**Highlights:**

- High resolution structures of the redox active cofactor TXNL1 in complex with the human 26S proteasome solved by time-resolved cryo-EM.
- TXNL1 binds the catalytic groove of the main proteasomal deubiquitinase Rpn11 and coordinates its active-site Zinc specifically in substrate-degrading states of the proteasome.
- Visualizing a partially unfolded intermediate of the mEos model substrate during processing.
- Structures of the actively degrading human proteasome reveal all spiral-staircase registers of the AAA+ ATPase hexamer with unexpected nucleotide occupancies that indicate asymmetric ATP hydrolysis mechanisms, conformational changes with burst phases, and thus new models for hand-over-hand substrate translocation.

## Introduction

The 26S proteasome unfolds and degrades hundreds of cellular proteins with highly diverse structural and chemical features to maintain protein homeostasis, remove misfolded, damaged, or obsolete regulatory proteins, and generate peptides with second-messenger functions ^1,2^. The proteasome consists of one or two 19S regulatory particles (RP) that bind to either end of the 20S core particle (CP), a barrel-shaped structure with an internal degradation chamber sequestering the proteolytic active sites of the proteasome ^3^. The RP includes the lid and base subcomplexes. The lid consists of the non-catalytic subunits Rpn3, 5-9, 12, and Sem1 (human PSMD3, 12, 11, 6, 7, 13, 8, and SEM1), as well as the Zn^2+^-dependent deubiquitinase (DUB) Rpn11 (human PSMD14). The base subcomplex contains the heterohexameric AAA+ motor with six distinct ATPase subunits in the order Rpt1, Rpt2, Rpt6, Rpt3, Rpt4, and Rpt5 ^4^ (human PSMC2, 1, 5, 4, 6, 3), the large scaffolding subunit Rpn2 (human PSMD1), and the three intrinsic ubiquitin receptors, Rpn1 (human PSMD2), Rpn10 (human PSMD4), and Rpn13 (hRpn13) ^5–8^. For simplicity we will use the yeast nomenclature of proteasomal subunits that is well established for structure-function studies in the field.

Each Rpt subunit includes a N-terminal helix that in the assembled hexamer forms a coiled-coil with one of the neighboring Rpts, a small domain with oligonucleotide/oligosaccharide-binding (OB) fold that forms an N-terminal domain ring (N-ring), and a C-terminal ATPase domain with a large and a small AAA+ subdomain that constitute the ATPase motor ring ^9^. The majority of proteasomal substrates are targeted for degradation through the enzymatic attachment of poly-ubiquitin chains, which bind to a ubiquitin receptor of the proteasome and allow an unstructured initiation region of the substrate to enter the central channel of the ATPase motor ^10–13^. A conserved loop structure, the so-called pore-1 loop, protrudes from each Rpt subunit into the central channel and sterically engages the substrate polypeptide ^14,15^. ATP-hydrolysis-driven conformational changes in the ATPase hexamer then apply mechanical force for substrate unfolding and translocation into the 20S CP. Rpn11 sits above the central pore of the ATPase motor and cleaves ubiquitin chains *en bloc* from translocating substrates ^16^.

Prior to substrate binding and engagement with the ATPase subunits, the 26S proteasome predominantly resides in a resting or engagement-competent state, in which Rpn11 is offset from a coaxial alignment with the N-ring, allowing substrates to enter the central channel ^3,17^. In this resting state, the N-ring, AAA+ motor ring, and the 20S CP are misaligned, and access to the degradation chamber is restricted by closed axial gates of the 20S CP. Pore-1 loop engagement with the substrate polypeptide induces major conformational changes of the 19S RP to processing states, which are characterized by a rotated lid subcomplex and a wider central channel with coaxially aligned N-ring, ATPase ring, and an open-gated 20S CP ^17^. Rpn11 shifts to a central position above the Rpt hexamer that facilitates substrate deubiquitination in a co-translational manner ^11,16,17^. Previous cryo-EM structural studies of the human and yeast 26S proteasomes in which substrate translocation was stalled either through addition of the non-hydrolysable ATP analog ATPγS or by inhibiting Rpn11-mediated deubiquitination, respectively, revealed the Rpt hexamer with different nucleotide occupancies and forming various right-handed spiral-staircase arrangements around the engaged substrate polypeptide ^18–20^.

Most of these stalled proteasome states showed 3 - 5 ATP-bound Rpt subunits at the top of the staircase and 1 – 3 ADP-bound or nucleotide-free subunits toward the bottom. 4 – 5 of the six subunits were engaged with the substrate polypeptide through their pore-1 loop, while 1-2 subunits were in an off or “seam” position, with usually one subunit located between the bottom and top subunits of the staircase. These observations led to a “hand-over-hand” translocation model, in which the second-to-last subunit in the spiral staircase hydrolyzes ATP and releases the phosphate, causing the neighboring ADP-bound bottom subunit to disengage from the substrate and move as the “seam” subunit to the top of the staircase. Subsequently this seam subunit exchanges ADP for ATP and re-engages the substrate, which pushes the other, already substrate-engaged subunits downward in the staircase by ∼ 5 Å and leads to a corresponding substrate translocation step of ∼ 2 amino acids ^18^. Hence, hydrolysis events appear to progress counterclockwise in the ATPase ring and phosphate release may drive the conformational changes that propel the substrate through the central channel.

Alternatively to ubiquitin modifications, substrates can be targeted to the proteasome through ubiquitin-like modifiers such as FAT10 (ubiquitin D, ^21^) and through interactions with proteasome-binding cofactors ^22–25^. FAT10 contains two ubiquitin-like (UBL) domains, one of which is bound in a partially unfolded state by the cofactor NUB1 that delivers it to the proteasome for motor engagement and subsequent degradation of FAT10 and any attached substrate moiety ^25^. In addition to these cofactors with substrate-delivery and chaperone functions, there is a multitude of proteasome-binding cofactors with enzymatic activities, including the deubiquitinases USP14/Ubp6 and UCH37, the ubiquitin ligase Hul5/Ube3c, the phosphatase UBLCP1 and the thioredoxin-like protein TXNL1 ^26–30^. Except for USP14/Ubp6, most of these enzymatic cofactors are only poorly understood regarding their proteasome interactions, mechanisms of substrate processing, and potential coordination with other proteasome functions. The redox-active TXNL1 is expressed in many cell-types and was previously proposed as a nearly stoichiometric component of the human 26S proteasome, whereby it uses its C-terminal PITH (proteasome interacting thioredoxin) domain (also known as DUF1000 domain) for binding, while potentially reducing disulfide bonds with its N-terminal thioredoxin (TRX) domain ^29,31^. However, the details of this interaction and its regulation remained elusive.

Here we used cryo-EM and *in vitro* biochemical studies on a reconstituted system to characterize the interactions of TXNL1 with resting and actively degrading human proteasomes. We found that TXNL1’s PITH domain binds on top of Rpn11, making contacts with Rpn10 and Rpn2, such that its catalytic TRX domain is positioned near the substrate entrance to the ATPase motor. TXNL1 thereby forms low-affinity interactions with the resting-state proteasome, where it is observed in multiple conformations yet similar locations above Rpn11. On actively degrading proteasomes, however, TXNL1 binds with high affinity through additional interactions of its C-terminal tail with the active-site Zn^2+^ and the catalytic groove of Rpn11 in a potentially deubiquitination-incompetent conformation. Extensive 3D classification of the substrate-degrading proteasome revealed at least six distinct states of the ATPase motor with previously unknown conformations and nucleotide occupancies that shine new light on the mechanisms of mechanochemical coupling and substrate translocation by the 26S proteasome and AAA+ motors in general. These states suggest burst-phase-like ATP hydrolysis that may aid in substrate unfolding and better explain existing discrepancies between single-molecule experiments and structure-based models for hand-over-hand substrate translocation. In addition, differential interactions of these motor states with TXNL1 indicate a sophisticated coordination of degradation steps, whereby TXNL1 does not interfere with initial or co-translocational substrate deubiquitination.

## Results

### High resolution structures of TXNL1 bound to the human 26S proteasome

We identified TXNL1 bound to endogenous proteasomes when we further processed a previously collected dataset of the human 26S proteasome during active degradation of a FAT10-fused Eos3.2 model substrate in the presence of the NUB1 cofactor ^25^. These affinity-purified proteasomes from HEK293 cells showed sub-stoichiometric amounts of TXNL1, whose binding is sensitive to the ionic strength the solution, such that proteasomes free of endogenous TXNL1 could be prepared by high-salt washes (Extended Figure 1A). To increase TXNL1 occupancy and the resolution of our cryo-EM structures, we saturated the human proteasomes with recombinant TXNL1 purified from *E. coli* (Extended Figure 1B). Robust redox activity of this recombinant TXNL1 was confirmed with an insulin reduction assay, in which resolving insulin’s disulfides leads to aggregation and a quantifiable increase in sample turbidity (Extended Figure 1C).

We used time-resolved cryo-EM to investigate the conformational landscape of the substrate-processing 26S proteasome 2 min after its incubation with excess of FAT10-Eos model substrate, NUB1 cofactor, and TXNL1 (Extended Figure 2). Particles were separated into two major groups, the resting state (RS, also known as the s1 state for yeast proteasomes) with ∼ 414K particles, and the processing states (PS, also referred to as non-s1 states) with ∼ 335K particles (Extended Figure 2). Deep 3D classification of the processing-state proteasomes resulted in at least 6 distinct conformations with global resolutions of 3 - 3.5 Å, showing the Rpt hexamer in distinct spiral-staircase registers (Extended Figure 2 and Extended Figure 3). We refer to these processing states based on the highest substrate-engaged Rpt subunit in the staircase as PS_Rpt1,_ PS_Rpt5_, PS_Rpt4,_ PS_Rpt3_, PS_Rpt6_ and PS_Rpt2_, represented by ∼19.3k, ∼112.2k, ∼29.5k, ∼32.2k, ∼24.2k and ∼73.1k particles, respectively (Extended Figure 2 and Extended Figure 3). All of them show the typical characteristics of the substrate-engaged proteasome, with a shifted and rotated lid subcomplex, the Rpt4/Rpt5 coil-coil positioned near Rpn10, a co-axial alignment of the ATPase ring with the open-gated 20S CP, and substrate polypeptide threaded through the central channel. We initially focus on the most abundant and highest resolution (∼3 Å) state, PS_Rpt5_ (Figure 1A, Extended Figure 3). In this state, all five subunits from Rtp5 at the top to Rpt3 at the bottom of the staircase are substrate-engaged through their pore-1 loop, while Rpt4 is the non-engaged seam subunit and at low resolution, likely due to a continuous motion from the bottom to the top position. Through 3D classification focused on the ATPase hexamer of PS_Rpt5_ we could resolve sub-states A and B that differ in the position of Rpt4 (Extended Figure 2), and we will concentrate our subsequent discussion of structural features on state A, PS_Rpt5A_.

**Figure 1:**
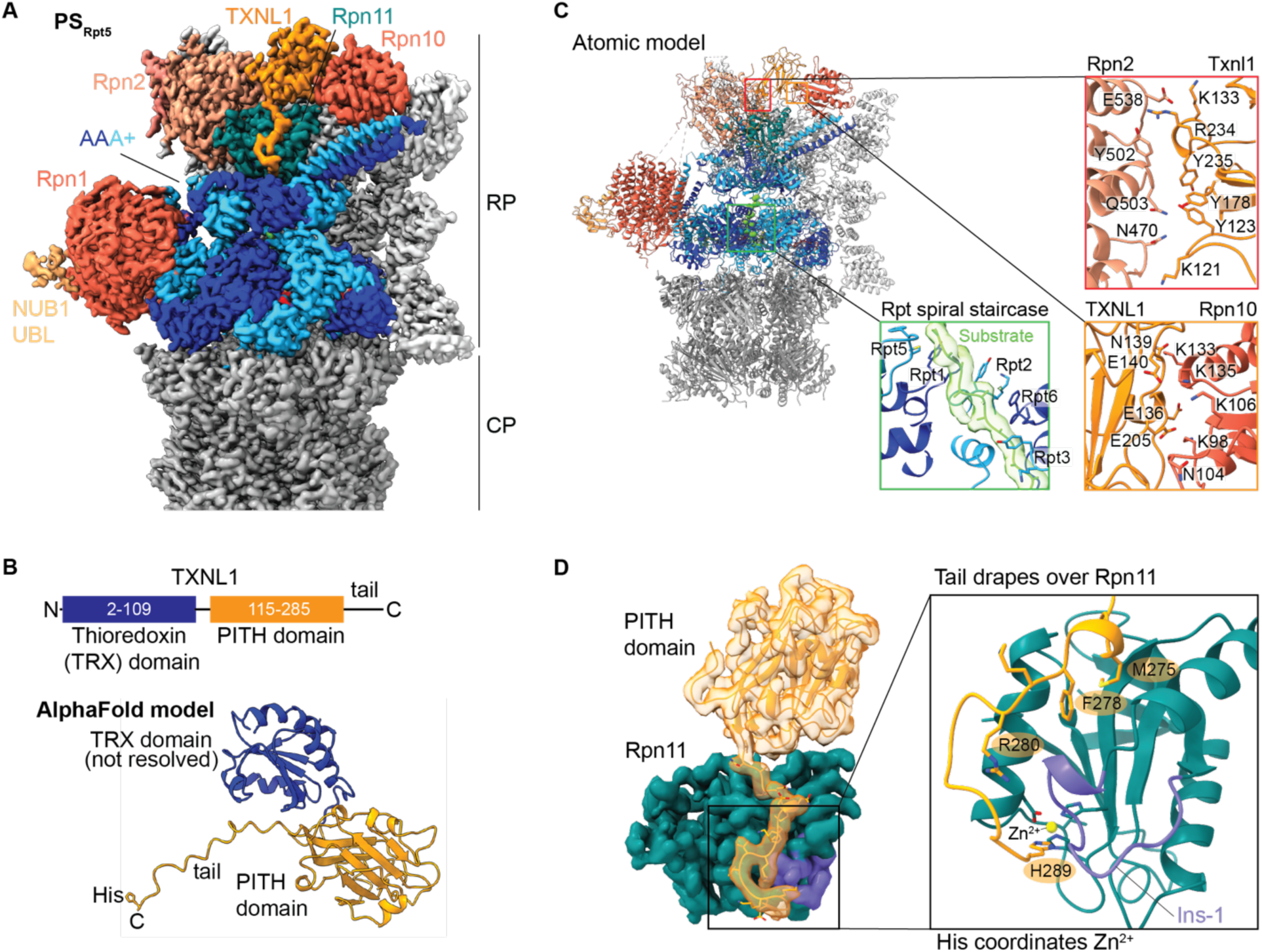
TXNL1 interacts with the substrate degrading human 26S proteasomes and co-ordinates the active site zinc of Rpn11. A) Density of TXNL1’s PITH domain bound to the human 26S proteasome in the processing state PS_Rpt5_ with Rpt5 at the top of the AAA+ ATPase spiral staircase and engaged with a translocating peptide during degradation of FAT10-Eos substrate delivered by NUB1. TXNL1’s PITH domain and C-terminal tail are shown in orange, the DUB Rpn11 in petrol, Rpn2 in salmon, Rpn1 and Rpn10’s VWA domain in tomato, the AAA+ ATPase hexamer in alternating dark blue and cyan, lid subunits (except Rpn11) in light grey, and the 20S CP in dark grey. B) Top: Schematic of TXNL1’s domain organization. Bottom: AlphaFold model of full length TXNL1. The N-terminal TRX domain is flexibly attached to the C-terminal PITH domain and therefore not resolved in our EM maps. C) Atomic model of PS_Rpt5_ with bound TXNL1. Zoom-in views on the right depict the interfaces of TXNL1 with Rpn2 and Rpn10, as well as the Rpt pore-1 loops forming a spiral staircase around the translocating substrate polypeptide. D) Left: Density for the PITH domain and its C-terminal tail bound to Rpn11, with Rpn11’s Ins-1 region shown in purple. Right: Zoom-in view of TXNL1’s C-terminal tail, with hydrophobic residues (M275 and F278) pointing toward Rpn11’s catalytic groove and the C-terminal H289 coordinating the active-site Zn^2+^.

The extra density above Rpn11 could be unambiguously assigned to the PITH domain of TXNL1 (Figure 1A,B). It contacts Rpn2 and the VWA (von Willebrand factor type A) domain of Rpn10 primarily through ionic interactions (Figure 1C), which explains the salt-sensitivity of this interaction. The N-terminal catalytic TRX domain of TXNL1 could not be resolved (Figure 1A), likely due to its flexible linkage to the PITH domain and consequent high mobility. Aligning AlphaFold models for the full-length TXNL1 with our structures places the TRX domain near the substrate entrance of the N-ring (Extended Figure 4C), and its mobility may aid in the reductive processing of a structurally diverse array of substrates. We observed the C-terminal tail of TXNL1 draped over Rpn11’s hydrophobic catalytic groove and the Insert-1 (Ins-1) loop that plays important role in binding the C-terminus of ubiquitin and regulating access to the deubiquitination active site (Figure 1D).

### TXNL1 coordinates the active site zinc of Rpn11 but does not inhibit deubiquitination

TXNL1’s interaction with the catalytic groove of Rpn11 is supported by hydrophobic interactions of the tail residues F278 and M275 (Figure 1D). Interestingly, the C-terminus of this tail reaches into Rpn11’s active site, where a conserved terminal histidine coordinates the catalytic Zn^2+^ (Figure 1D, Extended Figure 4A). The Ins-1 region of Rpn11 is a critical regulatory segment that switches its conformation depending on the state of the proteasome and the presence of ubiquitin. At least three distinct Ins-1 states have been described: an ‘open’ loop conformation in the resting-state proteasome, an active β-hairpin conformation in which Ins-1 forms a small beta sheet with the C-terminus of ubiquitin to position it for isopeptide cleavage during substrate deubiquitination, and a curled up ‘closed’ or inhibitory conformation typically found in substrate-processing states of the proteasome (Extended Figure 4B) ^16,18,19,32^. The specific conformation and interactions of TXNL1’s C-terminal tail is compatible only with the closed state of Ins-1, whereas a ubiquitin-bound state would lead to steric clashes, and the open state would be unable to form the necessary contacts. TXNL1’s tail may thus bind the catalytic groove of Rpn11 only on substrate-processing proteasomes when there is no deubiquitination. Previous structural and functional studies suggested that polyubiquitinated substrates first use a more distally located moiety in their ubiquitin chain to interact with a proteasomal ubiquitin receptor, before the proximal ubiquitin binds to Rpn11, followed by insertion of the substrate’s flexible initiation region into the central channel for motor engagement ^11,19,33^. This engagement triggers the conformational switch of the 19S RP from the resting to processing states and allows subsequent co-translocational deubiquitination ^11,34^. To test whether TXNL1 inhibits deubiquitination, we prepared a model substrate (Eos-I27^V15P^-tail) in which Eos3.2 was fused to the titin I27 domain with a destabilizing V15P mutation. The titin domain also carried a flexible C-terminal tail that contained a single lysine residue for Rsp5-mediated polyubiquitination and was long enough to allow degradation initiation. Measurements of proteasomal substrate degradation under multiple-turnover conditions, monitored through the decay of intrinsic Eos fluorescence, showed identical rates (0.54 +/- 0.06 min^-1^) in the absence and presence of excess TXNL1 (Extended Figure 1D). Similarly, monitoring substrate deubiquitination and peptide-product formation by SDS-PAGE with a N-terminally fluorescein-labeled Eos-I27^V15P^-tail substrate revealed no effect of TXNL1 on ubiquitin-dependent degradation (Extended Figure 1E), suggesting that TXNL1’s interaction with Rpn11 does not interfere with the removal of the affinity-conferring ubiquitin chain from a substrate upon motor engagement. All purified human proteasomes were treated with ubiquitin-propargylamine (PRG), a selective inhibitor of cysteine-dependent deubiquitination, which ruled out that other human proteasome-associated DUBs substituted for Rpn11 during ubiquitin-dependent degradation in the presence of TXNL1. Importantly, there was no indication for any *in vitro* degradation of TXNL1 (Extended Figure 1E), despite its location above the motor entrance and previous findings about endogenous TXNL1 being degraded upon arsenic treatment of mammalian cells ^31^.

### TXNL1 forms low-affinity interactions with resting-state proteasomes

Interestingly, although our cryo-EM data set showed a higher number of proteasome particles in the resting state, TXNL1’s tail was not detected interacting with Rpn11 in these proteasomes, where Rpn11’s Ins-1 loop is in the open conformation (Figure 2). Moreover, TXNL1’s PITH domain adopted an ensemble of orientations between Rpn2 and Rpn10 (Figure 2, Extended Figure 5, and Extended Figure 6). We classified particles into RS.1 and RS.2 states, which showed slight variations in the lid subcomplex, but no difference for the ATPase motor (Extended Figure 6). Local refinement of the PITH domain and 3D-variability analyses for both RS.1 and RS.2 allowed us to separate two major PITH-domain states that we term forward (49.8 % of particles) and backward (40.5% of particles) based on the PITH domain rotation (Figure 2A,B, Extended Figure 6), along with a smaller number of particles showing the PITH domain in intermediate positions. In the absence of C-terminal tail binding to Rpn11, the PITH domain thus appears to form ‘blurry’ interactions with the resting-state proteasome, using just a few contact points on Rpn2 and Rpn10’s VWA domain to swivel between forward and backward positions by ∼ 91 °. The forward position resembled the TXNL1 conformation we observed in the processing-state proteasomes, except for the C-terminal tail that did not contact Rpn11 and was therefore not resolved (Figure 2A, Figure 1A). Remarkably, the backward conformation utilizes overlapping, yet slightly different contact points on its PITH domain, Rpn2, and Rpn10, with again no involvement of the C-terminal tail (Figure 2C). These alternative orientations and their poorly resolved intermediate states thus provide an interesting example for “fuzzy” multi-mode interactions of a cofactor with a large protein complex that allow modulating affinity and function. 3D classification of a separate resting state of particles where Rpn1 is mobile and at lower resolutions also highlighted that forward and backward conformations were populated (Extended Figure 5).

**Figure 2:**
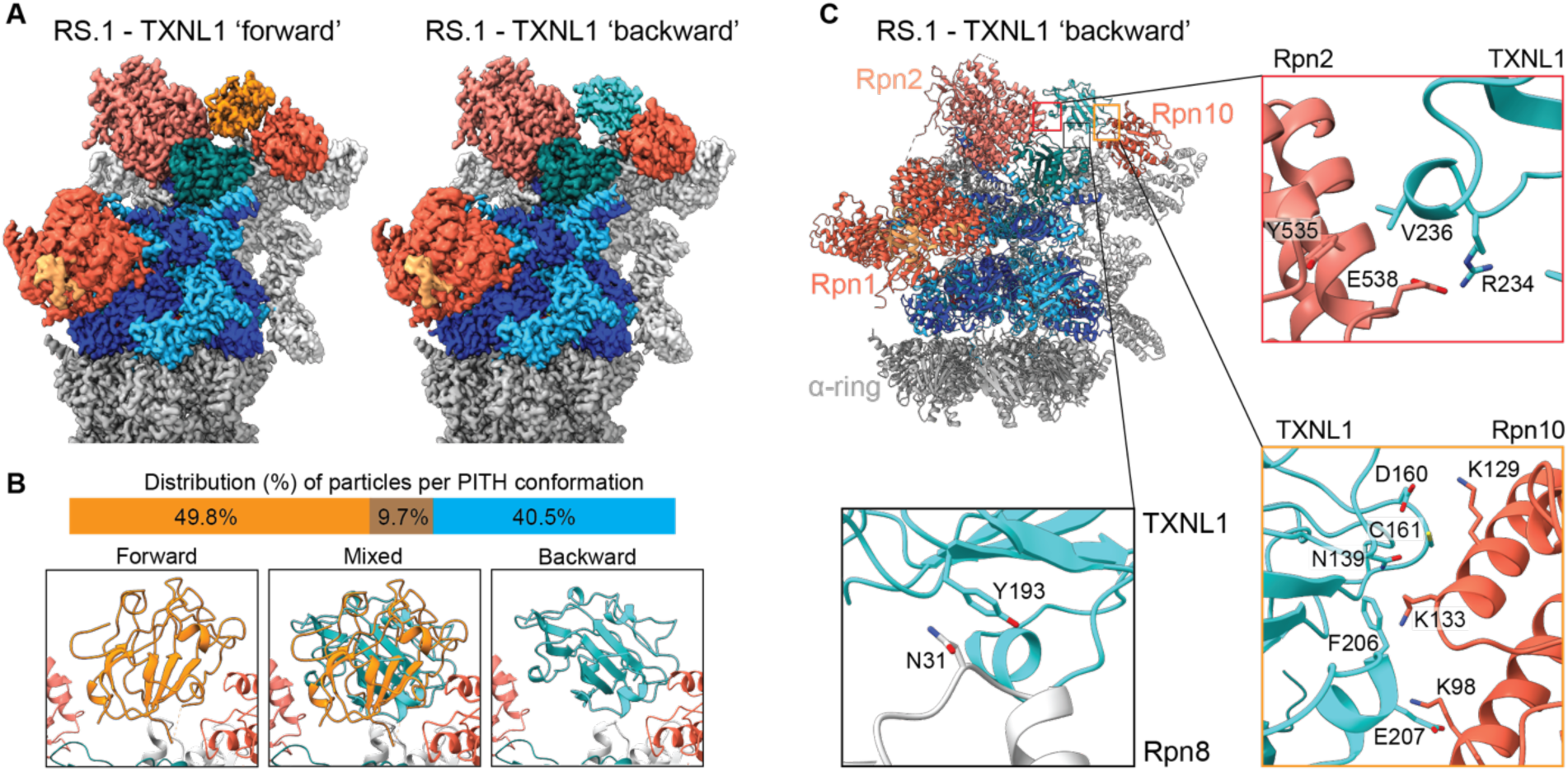
TXNL1 binds the resting-state proteasome in multiple conformations through contacts with Rpn2 and Rpn10’s VWA domain. A) Densities of the resting-state (RS.1) 26S proteasome bound to TXNL1’s PITH domain in the forward (left, PITH shown in orange) and backward (right, PITH shown in cyan) conformations. The forward position is identical to the location of the PITH domain on processing-state proteasomes, whereas the backward conformation is rotated by ∼ 90 °. RS.1 refers to a particular class of resting state proteasomes that differ from RS.2 through a slight shift in the regulatory particle (see Extended Figure 6). B) Particle distribution of resting-state proteasomes with TXNL1 bound in the forward, backward, or mixed conformations. Mixed conformations represent intermediate states due to PITH domain motions or particles with ambiguous probabilities of belonging to either conformation. C) Top left: Atomic model of the RS.1 proteasome with the PITH domain (cyan) is the backward conformation. Right and bottom: Zoom-in views of specific interactions between the PITH domain and Rpn2, Rpn10’s VWA domain, and Rpn8. The contacts with Rpn2 and Rpn10 strongly rely on ion pairs, while a polar-π interaction is at the center of the interface with Rpn8.

The backward conformation thereby appears to avoid any interference with deubiquitination upon substrate engagement. Indeed, for proteasomes that adopted a conformation with similarities to the previously reported E_A2_ or E_B_ states, which were proposed to reflect substrate deubiquitination ^19,20^, we observed TXNL1 in the backward conformation and, in addition, detected a low-resolution density bound to Rpn11 that resembles ubiquitin and could represent co-purified ubiquitin or a UBL domain of the FAT10 substrate (Extended Figure 5). Similar to the E_B_ state, these proteasomes show Rpn1 at a kinked angle relative to the ATPase ring, yet the ATPase ring has no translocating polypeptide in the central channel and is in a true resting-state conformation, like E_A2_. However, in contrast to the E_A2_ state that unexpectedly showed two ADPs bound to Rpt6 and Rpt5 across from each other in the hexamer, we observed five subunits bound to ATP and only the seam subunit Rpt6 in an ADP-bound state (Extended Figure 5). The backward conformation of TXNL1 is hence exclusively found in resting-state proteasomes, whereas the forward conformation is populated in both resting-state and actively degrading proteasomes, with the latter showing additional interactions of TXNL1’s C-terminal tail with Rpn11’s catalytic groove and active-site Zn^2+^.

### TXNL1 binds to actively degrading proteasomes with high affinity

Considering the structural findings described above, we wondered whether TXNL1 had a binding preference and formed higher-affinity interactions with actively degrading proteasomes. We therefore N-terminally labeled TXNL1 with fluoresceine-amidite (FAM) and measured its fluorescence polarization after mixing with excess human proteasome in either the absence or presence of FAT10 substrate and NUB1 cofactor. While the substrate-free, resting-state proteasome sample showed only low fluorescence polarization, actively FAT10-substrate degrading proteasomes caused a strong polarization increase (Figure 3A). Remarkably, this signal decreased as the substrate was consumed and the proteasome conformational equilibrium shifted back to the resting state, indicating the dissociation of TXNL1. Performing these polarization experiment with a TRX-domain deletion (PITH) or PITH-domain deletion (TRX) confirmed that this differential binding to human proteasomes is solely dictated by the Rpn11-interacting PITH domain (Extended Figure 7A). Furthermore, when using a 20S CP inhibitor to accumulate undigested FAT10 substrate in the degradation chamber and thereby stall substrate translocation through the 19S RP, we observed a constantly high polarization, indicative of TXNL1 tightly binding proteasomes that were trapped in the processing states (Extended Figure 7A). Titrating these stalled, substrate-engaged proteasomes allowed us to determine a *K_D_* of ∼ 35 nM for TXNL1 binding (Extended Figure 7B). By contrast, the TXNL1 titrations we performed with resting-state proteasomes for the cryo-EM structure determination let us estimate a *K_D_* in the tens of micromolar range, at least 2 orders of magnitude higher than for the actively degrading states. Hence, TXNL1 preferentially binds the substrate-processing human proteasomes with high affinity, whereas it switches to low-affinity “fuzzy” binding and dissociates when the proteasome is in the resting conformation.

**Figure 3:**
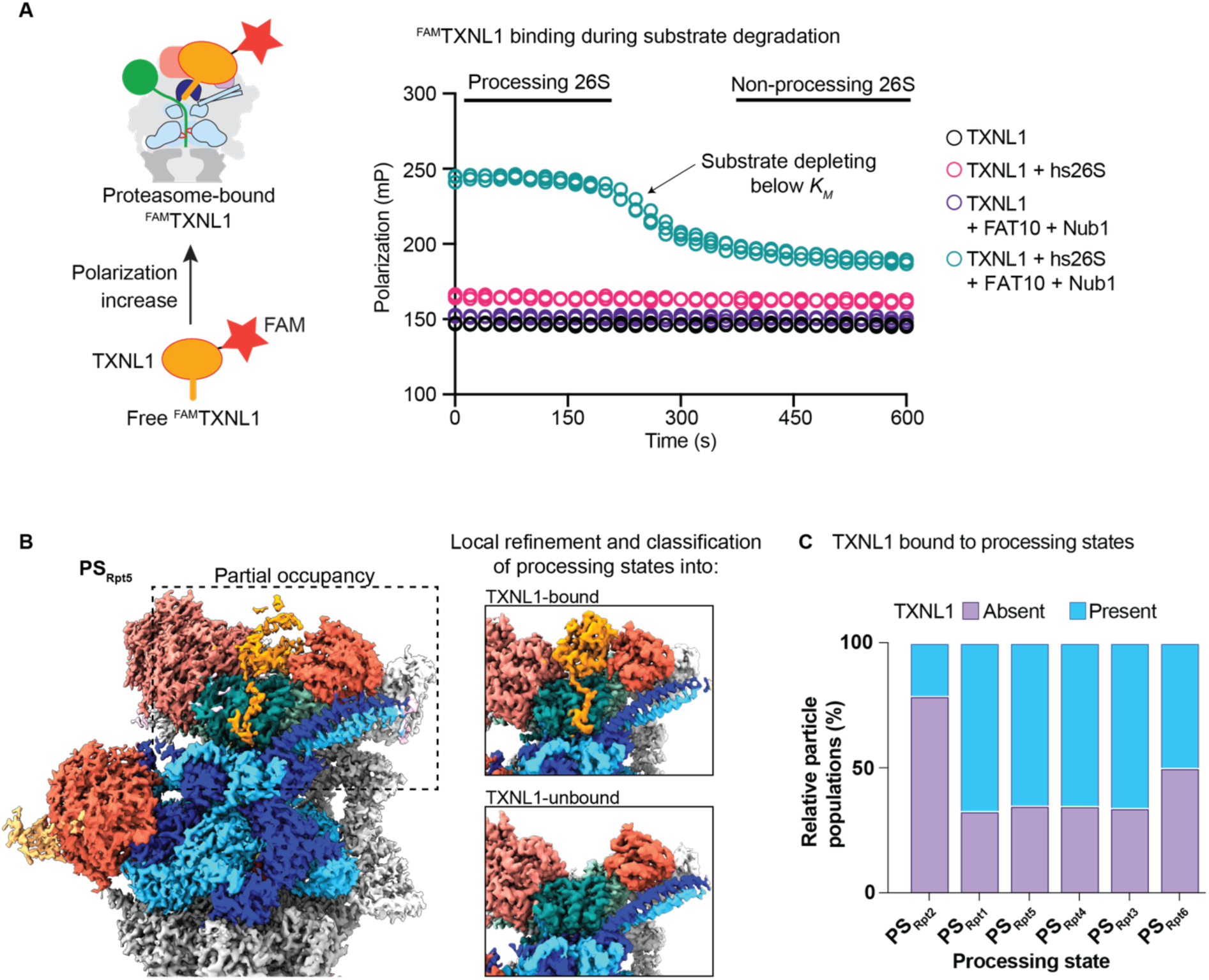
TXNL1 specifically interacts with processing-state 26S proteasomes. A) Proteasome binding and dissociation of N-terminally fluoresceine-amidite (FAM) labeled TXNL1 (50 nM) was measured by changes in fluorescence polarization after incubation with human 26S proteasome (500 nM) in the absence or presence of FAT10 substrate (10 μM) and the NUB1 cofactor (12 μM). B) Time-resolved cryo-EM of actively degrading proteasomes with sub-stoichiometric amounts of TXNL1 reveals TXNL1’s binding preference for processing states. Left: EM density of the proteasome in the PS_Rpt5_ conformation shows partial occupancy with TXNL1’s PITH domain (orange). Middle: Zoom-in views of example density maps from particles that were sorted into TXNL1-bound (top) and TXNL1-unbound (bottom) after local 3D classification and refinement of the PITH-domain density. C) Fractions of proteasome particles with and without bound TXNL1 as a function of the processing-state conformation.

### Time-resolved cryo-EM reveals conformation-specific TXNL1-binding preference

We further processed a previously collected dataset for the human 26S proteasome prepared 30 s after initiating the NUB1-mediated degradation of a FAT10-mEos3.2 substrate. Besides a large number of resting-state proteasome particles (77.3 %), we identified the substrate-degrading proteasome in the same six spiral-staircase ATPase states as observed for the 2 min time point (Extended Figure 8A). The resting-state proteasomes did not show PITH-domain density, consistent with the low concentration of the sub-stoichiometrically co-purified TXNL1 and its low-affinity binding to the resting state. For the processing-state proteasomes, we masked each Rpt subunit and used local refinements to improve the resolution of the ATPase hexamer for model building (Extended Figure 8A). The staircase states PS_Rpt1,_ PS_Rpt5_, PS_Rpt4,_ PS_Rpt3_, PS_Rpt6_ and PS_Rpt2_ were represented by ∼26.7k, ∼66.8k, ∼16.4k, ∼15.6k, ∼10.7k and ∼31.2k particles, respectively, and refined to moderate global resolutions of 3 – 4.5 Å (Figure 3B; Extended Figure 8A). There were no complete TXNL1 occupancies in these states, but alignments focused on the PITH domain allowed us to achieve high resolution (∼ 2.8 Å) and the unambiguous assignment of TXNL1 (Figure 3B; Extended Figure 8B,C). 3D classification of each PS conformation sorted particles into TXNL1-bound and TXNL1-unbound, and we noted a clear bias of PS_Rpt2_ for being TXNL1-free (Figure 3C, Extended Figure 8B). Interestingly, PS_Rpt2_ differs from the other substrate-processing states by showing a larger gap between the N-terminal coiled coil of Rpt4/Rpt5 and Rpn11’s Ins-1 region (Extended Figure 4D). This larger gap allows an Ins-1 conformation with possibly higher dynamics that may destabilize its interactions with the C-terminal tail of TXNL1. TXNL1 therefore binds PS_Rpt2_ similar to the resting-state proteasome, with low affinity and not involving its tail. It is tempting to speculate that the Ins-1 conformation in PS_Rpt2_ is less inhibitory for deubiquitination and more readily transitions to the active hairpin state that forms a beta sheet with the C-terminus of ubiquitin. PS_Rpt2_ may therefore represent a processing state that is suited for the removal of ubiquitin chains during substrate translocation or unfolding.

Overall, the structures of the degrading human 26S proteasome in the presence of sub-stoichiometric TXNL1 amounts further support our findings that TXNL1 forms high-affinity interactions only with the actively processing proteasome states, except for PS_Rpt2_, in which low-affinity binding may prevent TXNL1’s interference with co-translocational deubiquitination.

### Capturing an unfolding intermediate of the Eos substrate

Using the dataset for proteasomes with saturating TXNL1 at 2 min after initiating FAT10-substrate degradation, we visualized the PS_Rpt2_ conformation with an unfolding intermediate of the Eos substrate above the entrance to the ATPase channel and pulled against Rpn11 (Figure 4A). N-terminal of the Eos domain we were able to model the linker to FAT10’s C-terminus and continuously all amino acids up to FAT10’s residue 154, as they span through the N-ring and the ATPase spiral staircase (Figure 4A,B). The Eos domain is partially unfolded, with its FAT10-attached N-terminal β1 strand extracted from the beta barrel and the neighboring β2-β3 and β5-β6 hairpins strongly distorted (Figure 4C). Interestingly, β5 and β6 appear to maintain their hydrogen bonds as a hairpin, despite being pulled against Rpn11 and bent to parallel the extracted β1 strand going into the ATPase pore (Figure 4E,F). We were also able to model the Eos chromophore, a cyclic imidazole group formed from residues His227, Try228, and Gly229 (Figure 4D). Due to the displacement of β1, β5, and β6 from the barrel, the environment of the chromophore is strongly distorted (Figure 4E). It is therefore likely that Eos loses its fluorescence while forming this intermediate, whose subsequent further unfolding appears to be rate-limiting for the degradation of the FAT10-Eos substrate. Indeed, in previous biochemical studies we found that the Eos fluorescence during degradation under single-turnover conditions decayed with a time constant of *τ* = 18 s, whereas multiple-turnover degradation showed a time constant of *τ* = 50 s, likely determined by unfolding of the non-fluorescent intermediated ^25^. The Eos unfolding intermediate uses hydrophobic residues in β1, β5, and β6 that are normally part of the hydrophobic core to interact with the surface of Rpn11. In particular, M172, M274, and F279 engage in hydrophobic interactions with Rpn11’s Ins-1 region and V125 of the α3 helix (Figure 4F), thus stabilizing the intermediate in a particular position on Rpn11 that overlaps with the binding site of the TXNL1 C-terminal tail (Figure 4G). Future studies with other substrates will have to address whether this surface area of Rpn11 is specifically contacted only by Eos or more generally used to pull proteins against and potentially assist in their mechanical unfolding. Although PS_Rpt2_ is not the most abundant conformation of the substrate-degrading proteasome in our datasets (Extended Figure 2), it is the only one where an unfolding intermediate of the FAT10-Eos substrate could be visualized. Furthermore, the spiral staircase state with Rpt2 at the top was observed in previous structural studies of the yeast and human proteasomes with stalled substrates ^18,20^, suggesting that this register of the ATPase motor may play important roles during substrate unfolding or deubiquitination. Importantly, even PS_Rpt2_ proteasomes without the Eos unfolding intermediate showed a major reduction in TXNL1 binding (Figure 3C), indicating that high-affinity TXNL1 tail binding is disfavored by the PS_Rpt2_ conformation itself and not simply prevented by steric hindrance with the Eos unfolding intermediate pulled against Rpn11.

**Figure 4:**
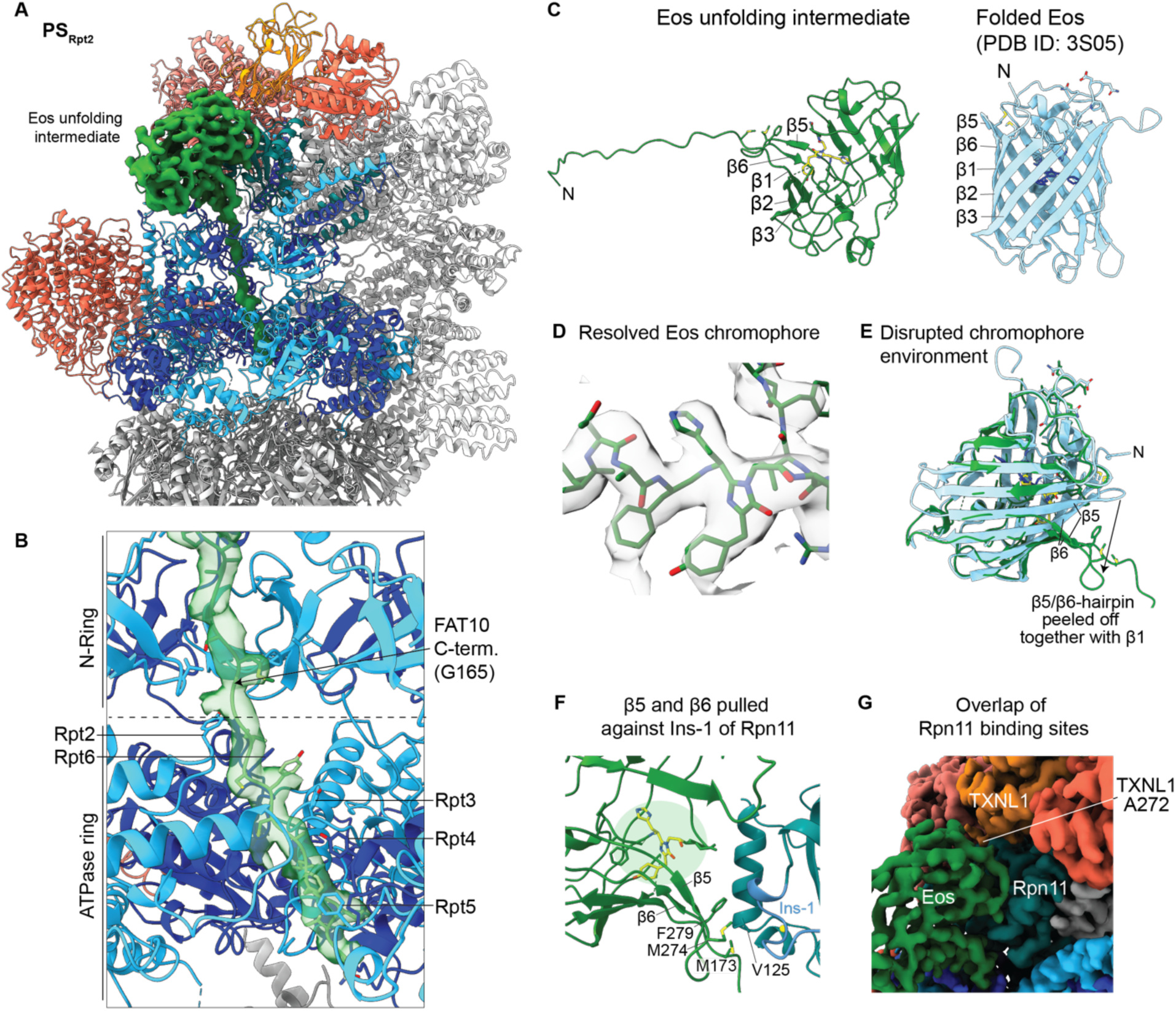
Visualizing a partially unfolded Eos intermediate during active degradation by the 26S proteasome. A) Atomic model of the human proteasome in the PS_Rpt2_ state during degradation of the FAT10-Eos model substrate. EM density (green) is shown for the substrate, with the partially unraveled beta-barrel of Eos pulled against Rpn11 and a translocating polypeptide spanning the central channel of the ATPase motor. B) Focus on the Rpt hexamer with the EM density and atomic model (green) shown for the translocating substrate, which is engaged by a staircase of ATPase domains. The substrate density is at high enough resolution to identify the C-terminal portion of FAT10 inside the ATPase ring. C) Comparison between the partially unfolded Eos intermediate (left, green) and the crystal structure of folded Eos (right, cyan; PDB ID: 3S05), with the chromophore shown in yellow and dark blue, respectively. The intermediate has β1 as well as the β2-β3 and β5-β6 hairpins partially pulled off from the beta barrel. D) EM density and atomic model for the chromophore in partially unfolded Eos. E) Overlay of the folded (cyan) and partially unfolded Eos (green) show the disruption of the chromophore environment that likely lead to a loss of fluorescence. F) Hydrophobic residues in β1 and the β5-β6 hairpin that normally face the Eos hydrophobic core interact with Rpn11 near the catalytic groove and the Ins-1 loop. G) EM density for proteasomes with bound TXNL1 and the Eos unfolding intermediate show an overlap of binding sites for Eos and TXNL1’s C-terminal tail on Rpn11.

### Asymmetric ATP hydrolysis in the Rpt hexamer

For both datasets of the human proteasome at 30 s and 2 min after substrate addition we observed non-substrate-engaged proteasomes in the typical resting-state conformation, with Rpt3 at the top of the staircase, Rpt2 at the bottom, and Rpt6 as the seam subunit bound to ADP, while all other subunits were ATP-bound. The substrate-engaged proteasomes in both datasets showed the six staircase conformations represented by vastly different particle populations (Figure 5, Extended Figure 9C). The resolutions were generally high enough to reliably determine the identities of the bound nucleotides and distinguish between ADP and ATP (Extended Figure 9B). For some of the more mobile seam subunits with lower resolution overall and for the nucleotide, we assumed ADP based on the subunit position in the staircase. PS_Rpt5_ was clearly the dominant state with particle numbers in the data sets ∼ 2 times higher than expected if all states were equally likely, and 4-5 times higher than three of the least populated motor states (Figure 5). Assuming that the numbers of particles in a particular conformation correlate with the lifetime of that state during ATP hydrolysis and the spiral-staircase progression in the Rpt hexamer, we postulate that the proteasomal ATPase motor functions by an asymmetric firing mechanism. This is further supported by varying distributions of nucleotide states, with processing states containing 4 ATPs and 2 ADPs, 3 ATPs and 3 ADPs, or only 2 ATPs and 4 ADPs (Figure 5). Assuming a counterclockwise progression of ATP hydrolysis, conformational changes, and nucleotide exchange events around the Rpt hexamer, we discuss the processing states in the corresponding counterclockwise order by which they may transition into each other. Although we describe discrete conformations for an almost complete cycle of ATP hydrolysis, it should be noted that we are likely missing rarer conformations and cannot assess continuous motions or very short-lived intermediates at high enough resolution for modeling, especially for the seam subunits.

**Figure 5:**
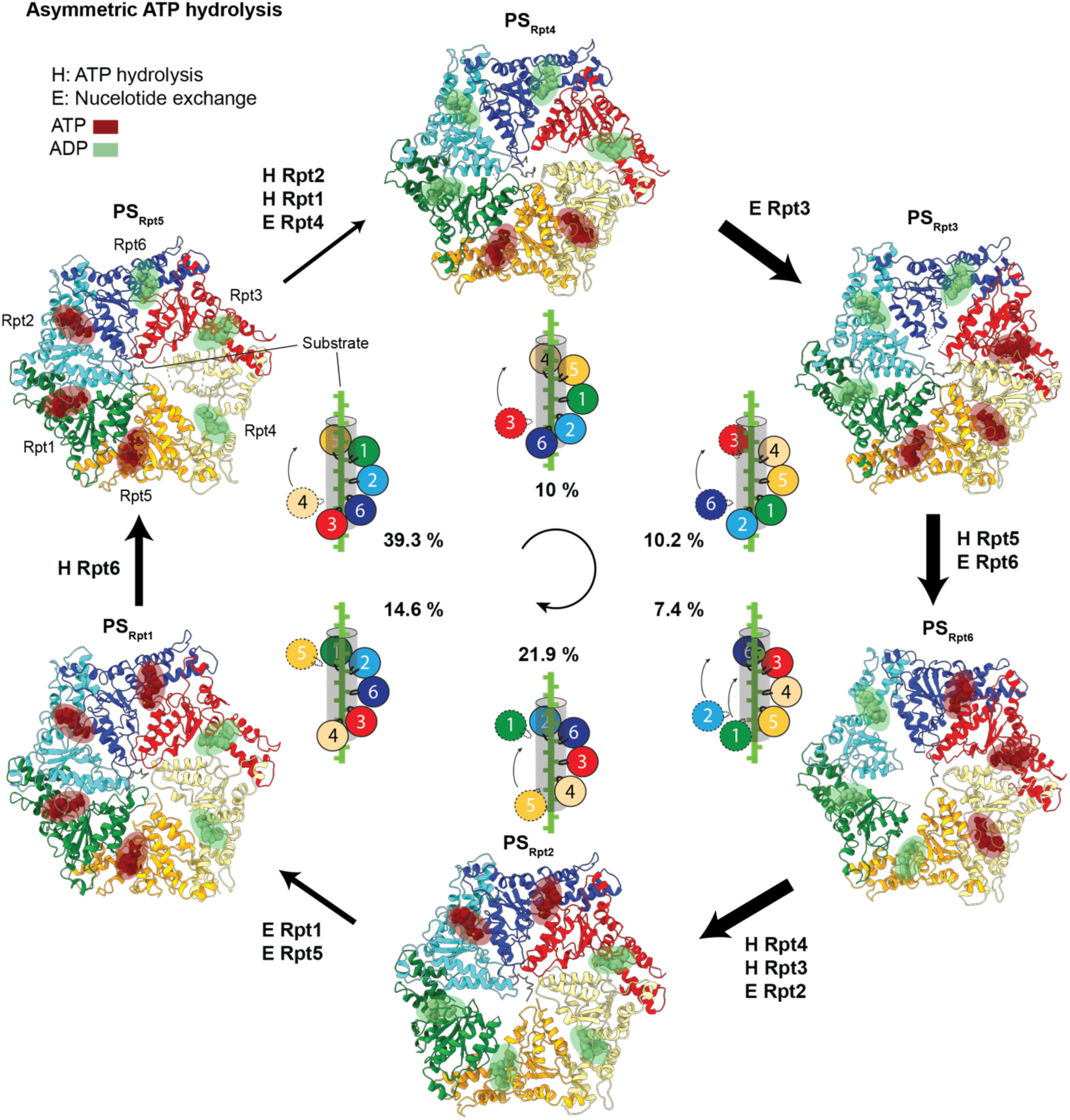
Conformational landscape of the proteasomal AAA+ motor with six distinct spiral-staircase registers during asymmetric ATP hydrolysis and active substrate processing. Atomic models show the AAA+ ATPase domains in different colors with the engaged substrate in the central channel, bound ATP depicted in red, and ADP in green. States are arranged in a cycle based on plausible order of events and transitions. The schematics depict the spiral-staircase arrangements and Rpt contacts with the substrate. Calculated cryo-EM particle distributions (in %) demonstrate asymmetries in the lifetime of individual processing states and the kinetics of their transitions, which is highlighted by different size arrows. For each transition the ATP-hydrolysis (H) and ADP-to-ATP exchange (E) events are indicated.

The most common conformation PS_Rpt5_ showed five subunits, Rpt5, Rpt1, Rpt2, Rpt6, and Rpt3, engaged with the substrate polypeptide, and Rpt4 as the disengaged seam subunit (Figure 5; Extended Figure 9A). At the top of the staircase Rpt5, Rpt1, and Rpt2 are ATP-bound, whereas Rpt6, Rpt3, and Rpt4 at the bottom have ADP in their active sites. Like for the 4D state of the substrate-engaged yeast proteasome ^18^, this would indicate that Rpt4 is the next subunit to move to the top of the staircase and exchange ADP for ATP. However, the 4D state and the E_D2.1_ state of the human proteasome ^20^ both had 4 ATP and 2 ADP bound to their Rpt subunits, suggesting that in PS_Rpt5_ an additional ATP-hydrolysis event had occurred in Rpt6 without being coupled to the movement or nucleotide exchange of Rpt4.

The PS_Rpt4_ state has not been previously observed in structural studies of the substrate-bound yeast or human 26S proteasomes. Interestingly, we detected only 2 ATPs, bound to Rpt4 and Rpt5, while the other 4 Rpt subunits were bound to ADP (Figure 5), which is a strong deviation from the conventional views of always having 4-5 ATP-bound and 1-2 ADP-bound or empty subunits. Again, this suggests that ATP hydrolysis can occur further upwards in the spiral staircase and ahead of the large movements for translocation and nucleotide exchange of the substrate-disengaged seam subunit. These findings likely indicate a burst phase in ATP hydrolysis, whereby additional hydrolysis events occur faster than the movement of seam subunits toward the top of the staircase, possibly because substrate translocation and staircase progression are held up by a folded substrate domain at the entrance of the ATPase motor. ADPs thus accumulate in up to 4 subunits, and their release may lead to several subunits moving from the bottom to the top of the staircase in rapid succession or even simultaneously. Our results challenge conventional models for hand-over-hand translocation that were based on structures of proteasomal or other AAA+ ATPase with stalled substrates and proposed a strict coupling of individual ATP-hydrolysis, subunit-movement, and nucleotide-exchange events ^18^. Interestingly, the preceding PS_Rpt5_ is the most abundant and likely most long-lived conformation of the hexamer, and its transition to PS_Rpt4_ is linked to two ATP hydrolysis events, suggesting that the movement, nucleotide exchange, and substrate engagement of Rpt4 are slow steps during substrate processing. Consistently, Rpt4 was previously identified as particularly important for substrate degradation ^14^, and our recent single-molecule studies revealed major unfolding defects and increased substrate release when Rpt4’s pore-1 loop was mutated ^35^. We thus propose that Rpt4 plays a critical important role for mechanical unfolding, possibly mediating a power stroke of the Rpt hexamer.

The transition to the subsequent PS_Rpt3_ state involves Rpt3 moving to the top of the staircase and exchanging ADP for ATP. As there are no additional hydrolysis or exchange events, this leads to a hexamer with 3 ATPs and 3 ADPs (Figure 5). Due to the lower resolution (∼3.5 - 4Å), we cannot interpret the side-chain orientations for Rpt3’s pore-1 loop, yet the backbone appears positioned close to the substrate and ready to engage (Extended Figure 9A,D). Consequently, two subunits, Rpt6 and Rpt3, are disengaged, while the other four contact the substrate through their pore-1 loop.

The following PS_Rpt6_ state also showed 3 ATPs and 3 ADPs bound to the ATPase hexamer, meaning that during the transition from PS_Rpt3_ one hydrolysis event occurred in Rpt5, while Rpt6 exchanged ADP for ATP. Interestingly, there are visible gaps at all ADP-bound subunit interfaces, Rpt5/Rpt1, Rpt1/Rpt2, and Rpt2/Rpt6, and Rpn1 is highly mobile, likely because its anchoring Rpt1 and Rpt2 subunits are at the staircase seam and simultaneously disengaged from the substrate (Extended Figure 9A,D).

In the subsequent PS_Rpt2_ state, Rpt2 and Rpt6 are ATP-bound, while the other four subunits are bound to ADP, meaning that two hydrolysis events and one nucleotide exchange occurred during the transition from PS_Rpt6_. Rpt2, Rpt6, Rpt3, and Rpt4 are in contact with the substrate, whereas Rpt5 and Rpt1 are disengaged (Figure 5). Interestingly, Rpt1 is rotated up towards the top of the staircase (at level with Rpt2), with its pore-1 loop residue Y249 at ∼ 20 Å distance from the substrate and potentially sequestered by interaction with Rpt2 (Extended Figure 9A,D). Hence, during the transition from PS_Rpt6_, Rpt2 and Rpt1 moved together to the top, possibly due to their coupling by the associated Rpn1 subunit. Rpt5 at the bottom of the staircase is disengaged, yet very close to the substrate (∼ 8 Å, Extended Figure 9A,D). A similar staircase conformation with disengaged Rpt5 and Rpt1, referred to as 1D*, was observed for the yeast proteasome ^18^, yet assumed to be off pathway. Here we detected three states, PS_Rpt3_, PS_Rpt6_, and PS_Rpt2_, with two disengaged subunits, and similar staircase states, like E_D0_ and E_D1_ ^20^, were previously reported for the human proteasome. Transitions with two disengaged seam subunits may therefore be common or even an important principle for substrate translocation that includes bursts of hydrolysis and Rpt-subunit movements.

In PS_Rpt1_, we again observed five subunits, Rpt1, Rpt2, Rpt6, Rpt3 and Rpt4, in contact with the substrate and Rpt5 as the seam subunit, albeit at the top of the spiral staircase, ATP-bound, and ready to engage (Figure 5, Extended Figure 9A,D). This state thus immediately precedes PS_Rpt5_. There are four subunits, Rpt5, Rpt1, Rpt2, and Rpt6, bound to ATP, with Rpt3 and Rpt4 ADP-bound. PS_Rpt1_ is therefore directly comparable to the previously reported human E_D2.0_ ^20^ and yeast 5T ^18^ states.

## Discussion

Our cryo-EM structural and biochemical studies revealed how the redox-active TXNL1 interacts with the human 26S proteasome in a highly conformation-specific manner, which provides exciting new insights not only into the coordination of proteasomal activities and cofactor functions, but also the operating principles of the AAA+ ATPase motor.

Transient ionic interactions of TXNL1’s PITH domain with the Rpn2 and Rpn10 subunits of the resting-state proteasome lead to low-affinity binding in two orientations above the Rpn11 deubiquitinase and allow dynamic association and dissociation. By contrast, when the proteasome is in actively degrading conformational states, TXNL1’s PITH domain binds with high, nanomolar affinity through additional interactions of its C-terminal tail with Rpn11’s catalytic groove and a coordination of the active-site Zn^2+^. TXNL1 thereby takes advantage of the conformational switch in Rpn11’s Ins-1 region, which controls substrate deubiquitination and inversely coordinates it with TXNL1 binding to the proteasome. In the resting, substrate-free proteasome, the Ins-1 region is in an open loop state that facilitates ubiquitin binding and cleavage, yet disfavors the interaction with TXNL1’s C-terminal tail. Upon substrate engagement and proteasome transition to actively degrading conformations, the Ins-1 region switches to a closed loop state that interferes with deubiquitination, but facilitates TXNL1 tail binding and Zn^2+^ coordination. Interestingly, this holds true for all degrading conformations and spiral-staircase registers of the AAA+ ATPase motor, except for PS_Rpt2_, in which a slightly larger distance between Rpn11 and the neighboring Rpt4/Rpt5 coiled coil may allow an alternative, more relaxed Ins-1 loop conformation that disfavors TXNL1 tail binding and likely accommodates deubiquitination. We determined these TXNL1-bound proteasome structures during ubiquitin-independent, NUB1-mediated degradation of a FAT10 substrate, yet in combination with previous structural and functional studies of ubiquitin-dependent degradation our results suggest a compelling model for the coordinated proteasome binding of TXNL1 that does not interfere with deubiquitination, despite utilizing critical catalytic features of Rpn11 (Figure 6). After substrate recruitment to the resting-state proteasome, a proximal ubiquitin moiety of the affinity-conferring ubiquitin chain appears to bind to Rpn11 ^19,33^. TXNL1 does not interfere with this ubiquitin binding to Rpn11, as it only loosely interacts in this state and its tail does not contact the Ins-1 region. Insertion of the substrate’s flexible initiation region and engagement by the ATPase motor triggers the proteasome conformational switch, the onset of translocation, and the immediately subsequent or even simultaneous ubiquitin cleavage ^11,33,34^, after which TXNL1 can bind Rpn11 with high affinity. In case a substrate contains additional ubiquitin chains, the proteasome would not have to switch back to the resting state for their removal, as the PS_Rpt2_ spiral staircase would allow a co-translocational deubiquitination while TXNL1 is dissociated or transiently shifted to a low-affinity backward position on Rpn11 (Figure 6B). Thus, the conformationally selective binding of TXNL1 allows its presence near the motor entrance during mechanical substrate processing, yet prevents the interference with initial or co-translocational deubiquitination events, which is also supported by our ubiquitin-dependent degradation experiments that showed no inhibition by excess amounts of TXNL1.

**Figure 6:**
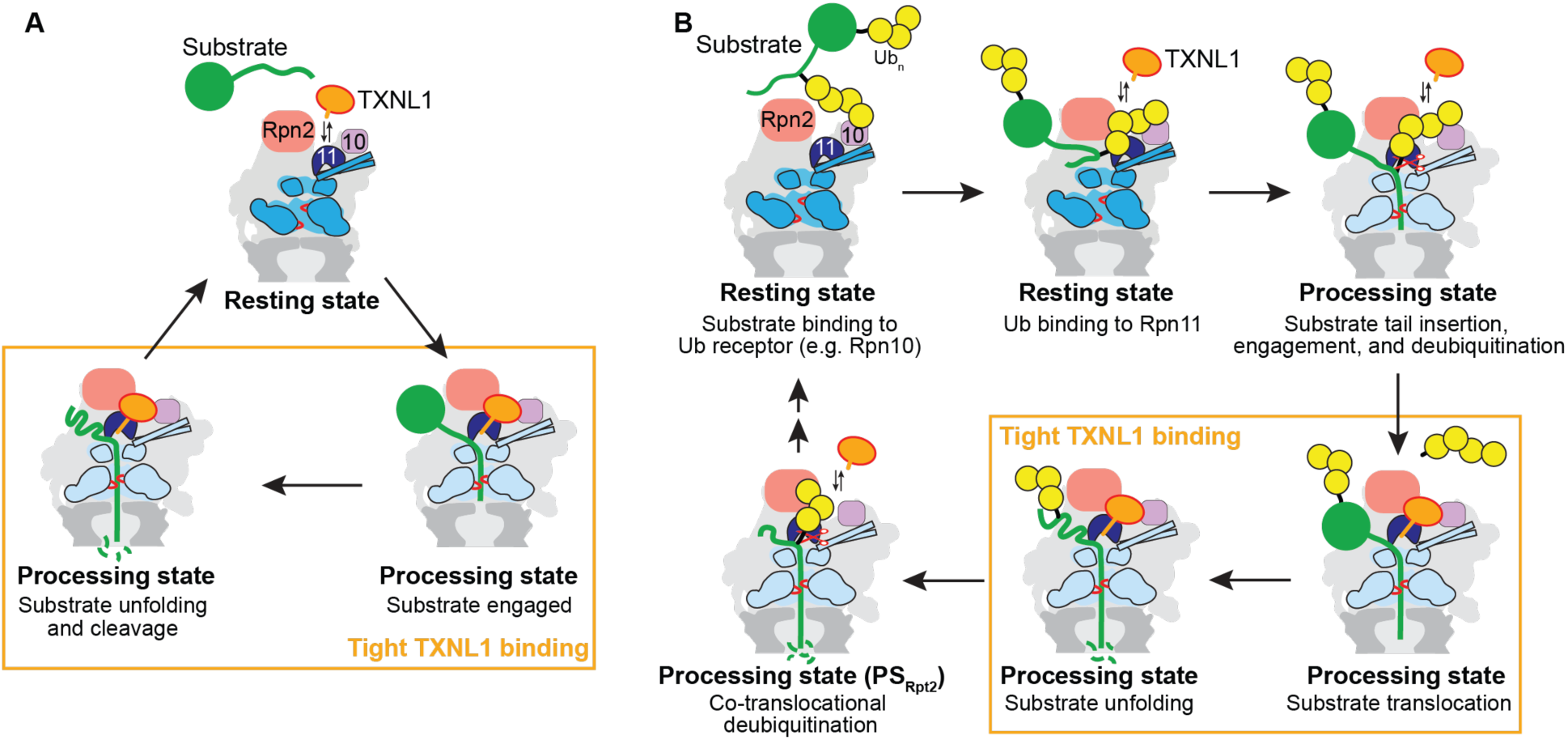
Model for the coordination of proteasome conformational changes, deubiquitination, and TXNL1 binding. A) TXNL1 loosely interacts with Rpn2, Rpn10, and Rpn11 of the resting-state proteasome and tightly binds processing states during substrate unfolding and translocation through additional contacts of its C-terminal tail with Rpn11’s catalytic groove, allowing a potential do-degradational reduction of oxidized substrates. B) The conformationally selective binding of TXNL1 prevents interference with deubiquitination during the turnover of ubiquitin-tagged substrates. After ubiquitin binding to a receptor and Rpn11 in the resting-state proteasome, the insertion of the substrate’s flexible initiation region into the ATPase motor induces the conformational switch to processing states and subsequent or simultaneous cleavage of the affinity-conferring ubiquitin chain by Rpn11. TXNL1 then binds with high affinity during substrate unfolding and translocation, yet lower TXNL1 affinity and the exclusion of its C-terminal tail from Rpn11 in the PS_Rpt2_ state would allow a co-translocational removal of any additional ubiquitin chains.

Although it cannot be ruled out that TXNL1 reduces proteasomal cysteines, it more likely reduces substrates during their unfolding and translocation, given the predicted position of the catalytic TRX domain near the entrance to the proteasomal central channel. Future studies will have to address the exact role of TXNL1 in the proteasome context under normal physiological or certain stress conditions. Severe oxidative stress that may lead to a prevalence of oxidized or crosslinked substrates was reported to cause the dissociation of the 26S proteasome, and the isolated 20S CP, rather than the 26S proteasome, is assumed to fulfil the task of damaged or aggregated protein degradation ^36^. Furthermore, TXNL1 levels were observed to decrease after arsenic treatment of mammalian cells ^31^, suggesting that TXNL1 is less of a stress regulator, but playing roles under normal physiological conditions. Indeed, knockdown of TXNL1 leads to a moderate accumulation of polyubiquitinated substrates ^29^. Although the cytosol is an overall reducing environment, the release of oxidizing compounds from peroxisomes, mitochondria, or the endoplasmic reticulum may cause local differences in the redox state that require the action of a thioredoxin-like protein for efficient proteasomal degradation. It was previously shown that the 26S proteasome can process unstructured polypeptides that are linked by two or even three disulfide bonds ^37^. However, degradation may be inhibited if disulfides prevent mechanical substrate unfolding by stabilizing individual domains or crosslinking several proteins, and TXNL1 may help resolve these crosslinks in a co-degradational manner.

Our structures of the TXNL1-bound, substrate-degrading proteasome also gave important new insights into the mechanisms for ATP-dependent unfolding and translocation by the AAA+ ATPase motor. For the first time we were able to observe all spiral-staircase registers of the substrate-processing hexamer, likely because we did not inhibit ATP hydrolysis or stall translocation as in previous studies. Interestingly, these hexamer states show 2 to 4 ATP-bound subunits, with the others being ADP-bound. The substrate-disengaged seam subunits are never detected in an empty state, and the ADP-to-ATP exchange therefore appears to occur rapidly after a seam subunit moved from the bottom to the top of the staircase. Individual transitions between staircase states involve single ATP hydrolysis and nucleotide-exchange events, but also double hydrolysis events, single hydrolysis without exchange, and single or double exchange events without hydrolysis, with either one or two seam subunits moving from the bottom to the top of the staircase (Figure 5). This significantly deviates from the existing hand-over-hand translocation model, which assumes that each translocation step depends on the coupled occurrence of one ATP hydrolysis event in a subunit near the bottom of the staircase, one seam subunit disengaging from the substrate and moving to the top, and one nucleotide exchange by that seam subunit before substrate re-engagement, leading to a downward movement of the polypeptide by ∼ 5 Å or 2 amino acids ^18^. Our findings indicate that ATP hydrolysis and phosphate release do not immediately drive the seam subunit’s disengagement from substrate. Instead, they possibly built up strain in the hexamer, especially for the transitions from PS_Rpt5_ to PS_Rpt4_ or PS_Rpt6_ to PS_Rpt2_ that involve two ATP-hydrolysis events. PS_Rpt5_ appears to be 4 - 5 times longer lived than the subsequent 3 states, and its seam subunit Rpt4 was shown in biochemical and single-molecule studies to be particularly important for efficient substrate unfolding ^14,35^, suggesting that the transition from PS_Rpt5_ to PS_Rpt4_ drives an unfolding power stroke of the motor. Interestingly, after the hexamer subsequently progresses rapidly through PS_Rpt4_, PS_Rpt3_, and PS_Rpt6_, it spends more time in the PS_Rpt2_ state, for which we observed a defined unfolding intermediate of the Eos3.2 model substrate, with a disordered fluorophore environment and the N-terminal beta strand extracted from the beta barrel. The progression through the entire ring cycle is thus highly asymmetric. 75 % of the time is spent by the Rpt hexamer transitioning through PS_Rpt2_, PS_Rpt1_, and PS_Rpt5_, which involves the movement of just two seam subunits, Rpt5 and Rpt4, to the top of the staircase (Figure 5). The presence of bursts, with a more rapid progression through certain spiral-staircase states, two ATP-hydrolysis events occurring in succession, or two seam subunit moving together, may thereby facilitate the mechanical unfolding of substrate proteins. Despite this plasticity in hydrolysis, nucleotide exchange, and subunit movement, the fundamental translocation step size appears to be ∼ 5 Å or 2 amino acids, driven by substrate engagement of one seam subunit at a time, which is linked, albeit not always directly, to one ATP-hydrolysis event in the ring.

Based on single-molecule optical tweezers studies of the related, homo-hexameric ClpX ATPase, we and others had previously proposed that these motors can take significantly larger steps ^38–42^, which seemed to contradict the structure-derived models with 2 amino acids per ATP. However, limitations in the spatio-temporal resolution of those measurements may have prevented the detection of smaller sub-steps. Indeed, recent single-molecule nanopore readings of polypeptide translocation by ClpX revealed a 2-residue step size ^43^, with overall translocation velocities similar to those observed in previous optical tweezers experiments.

In summary, our findings suggest that proteasomal ATPase subunits, and potentially other AAA+ motors, take a stochastic approach to substrate unfolding, using fast and slow ATP hydrolysis and exchange kinetics. Together with the slipping of pore loops that prevents motor seizing when encountering major obstacles ^34,35^, this mechanism allows repetitive unfolding attempts and likely higher probabilities in overcoming the diverse thermodynamic and kinetic barriers associated with the vast substrate pool of the 26S proteasome. The conformational asymmetries we detected in our study are overall consistent with those observed previously for the DUB-inhibited, substrate-engaged yeast proteasome ^18^ or the Usp14-bound human proteasome during ATPγS-inhibited Sic1-substrate degradation ^20^, where the PS_Rpt5_-equivalent states 5T and E_D2_ dominated the particle distributions. Apparent differences in these structures may have been caused by the type of substrate delivery or the processing step that proteasomes were primarily dealing with when captured. Hence, more proteasome structures in the presence of different substrates and cofactors will be necessary in the future to build a complete repertoire of motor states, further advance our understanding of ATP-hydrolysis-driven degradation, and elucidate the sophisticated coordination with enzymatically active cofactors.

## Methods

### Cloning

TXNL1 (Trp32) was gene synthesized with the following amino acid sequence that was codon-optimized for *E. coli* expression. TXNL1 DNA was ligated into an expression vector with an N-terminal His-TEV-FLAG (MGSSHHHHHHSSGENLYFQGHMDYKDDDDK), so that a GHMDYKDDDDK overhang was left on the N-terminus of TXNL1. All TXNL1 mutants and truncations were generated by site-directed mutagenesis with Q5 polymerase, phosphorylation, and ligation.

TXNL1 amino acid sequence MVGVKPVGSDPDFQPELSGAGSRLAVVKFTMRGCGPCLRIAPAFSSMSNKYPQAVFLEVDVHQCQGTAATNNISATPTFLFFRNKVRIDQYQGADAVGLEEKIKQHLENDPGSNEDTDIPKGYMDLMPFINKAGCECLNESDEHGFDNCLRKDTTFLESDCDEQLLITVAFNQPVKLYSMKFQGPDNGQGPKYVKIFINLPRSMDFEEAERSEPTQALELTEDDIKEDGIVPLRYVKFQNVNSVTIFVQSNQGEEETTRISYFTFIGTPVQATNMNDFKRVVGKKGESH

The mEos3.2-titin-tail construct was generated by Gibson assembly of His-TEV mEos3.2 before titin(V15P)-tail-intein-chitin binding domain (CBD). The titin-tail has one lysine followed by a PPPY motif for Rps5 ubiquitination. The P141C, C171A mEos-titin-tail construct was generated by site directed mutagenesis.

### Purification of TXNL1

TXNL1 was expressed in *E.coli* BL21* grown in TB medium by induction with IPTG (250 μM) when cells reached OD_600nm_ 0.6 - 0.8, followed by shaking (200 rpm) overnight at 16 °C. *E.coli* cells (from ∼2 L) were resuspended in 30 mL of lysis buffer (60 mM HEPES pH 7.4, 500 mM NaCl, 5% glycerol, 0.5 mM TCEP, 20 mM imidazole) with freshly added benzonase, before sonication and clarification to remove insoluble matter. The supernatant was then passed through Ni-NTA resin (∼2 mL) by gravity at room temperature (for ∼ 20 mins), before extensive washing to remove contaminants (10 column volumes, sequentially with lysis buffer). TXNL1 was eluted with lysis buffer supplemented with 250 mM imidazole (∼ 10 mL). Protease (His-TEV, 250 μg) was added and TXNL1 was dialyzed into 60 mM HEPES pH 7.4, 200 mM NaCl, 5% glycerol, 0.5 mM TCEP at room temperature for 2 hours, followed by overnight dialysis in fresh buffer at 4°C. Dialyzed TXNL1 was flown over equilibrated Ni-NTA resin and the flow through containing cleaved TXNL1 is collected and concentrated for gel filtration using an SD75 16/600 at 4°C. Protein is stored at -80°C in 60 mM HEPES pH7.4, 150 mM NaCl, 25 mM KCl, 10 mM MgCl, 0.5 mM TCEP as single use aliquots (typically ∼350 μM). Protein concentration is estimated using 280 nm.

### hs26S proteasome purification

As previously described, human 26S proteasome were purified from HTBH-Rpn11 tagged human HEK293 adapted to suspension culture ^25,44^. HEK293 cells were grown at 8 % CO_2_, 37 °C, and 120 rpm shaking in FreeStyle™ 293 Expression Medium with 2% (v/v) FBS. Cells were passaged twice a week at 5 × 10^5^ and newly thawed cells were grown with puromycin. Cell pellets from 4 L of culture were resuspended in 60 mM HEPES pH 7.4, 25 mM NaCl, 25 mM KCl, 5% glycerol, 10 mM MgCl_2_, 5 mM ATP, and 0.5 mM TCEP, supplemented with benzonase and EDTA-free proteasome inhibitor tablets. After lysis by a dounce homogenizer (usually 15X), lysate was sonicated on ice with low amplitude (20%), clarified for 60 min at 4°C, and flowed over pre-equilibrated Pierce™ High-Capacity Streptavidin Agarose (3 mL of resin). For preparing low salt treated hs26S proteasomes, resin was sequentially washed with lysis buffer 5X, whereas for high salt treated hs26S, resin was washed 4X with lysis buffer supplemented with 300 mM NaCl and 1X with lysis buffer without extra NaCl. Protein was eluted by TEV protease cleavage for 60 mins at room temperature and gentle agitation followed by concentration of the elution to ∼ 250 μL. Samples were clarified at 21,000 x g for 15 min before fractionation using an S6 increase 10/300 column in lysis buffer. Fractions containing 26S proteasome were concentrated, and concentrations were estimated in a Bradford assay with BSA as a standard and assuming a molecular weight of 2.6 MDa for the proteasome. Typically, 10 μL ∼ 4 μM aliquots were snap frozen in liquid N_2_ and stored at -80°C.

### Purification of Eos-Titin-tail substrate

Substrate and mutants were expressed as a His-TEV-Eos-titin I27(V15P)-tail-intein-CBD fusion in *E. coli* BL21* by growing cells at 37 °C to OD_600nm_ of 0.6, before cooling cells to 16 °C for ∼1 hour and induction with IPTG at 0.25 μM. Cell were harvested after 18 hours of induction, followed by lysis with sonication (12 cycles of 70% amplitude, 10 s on and 45 s off on ice) in 50 mM HEPES, 300 mM NaCl, 5% glycerol, 10 mM MgCl_2_, 20 mM imidazole with freshly added EDTA-free proteasome inhibitor tablets and benzonase. Lysate was clarified before flowing over Ni-NTA resin by gravity. Ni-NTA resin was washed extensively with lysis buffer without inhibitor tablets and benzonase. Protein was eluted with lysis buffer supplemented with 250 mM imidazole and bound to chitin resin, before washing with 5 column volumes of lysis buffer. Resin was suspended in lysis buffer overnight with the addition of 50 mM DTT and His-TEV protease. Columns were moved to room temperature for one hour, before collecting elution and flowing over freshly equilibrated Ni-NTA resin to remove uncleaved protein and His-TEV protease. The cleaned elution was concentrated using Amicon concentrator (30 kDa cutoff) before fractionation using an SD200 16/600 column in gel-filtration buffer (60 mM HEPES pH 7.4, 25 mM NaCl, 25 mM KCl, 5% glycerol, 10 mM MgCl_2_, 0.5 mM TCEP). After concentrating to ∼ 10 mg/mL, protein was snap frozen liquid N_2_ as single-use aliquots and stored at -80°C

### Additional proteins previously established

Fat10, NUB1, Sortase, TEV protease, 3C protease, His-ULP1 protease, Ubiquitin, Rsp5, His-TEV-mE1, Ubch7 and Ub-intein were expressed and purified using previously established protocols ^11,25^. Ubiquitin-Propargylamide (Ub-prg) was purified as previously described ^25^.

### Western blot

Samples were separated by SDS-PAGE and transferred to PVDF membranes (Thermo Scientific), before blocking in 5% milk TBS-T. TXNL1 primary antibodies (15289-1-AP, Thermo Scientific) were incubated at room temperature with the membranes for 90 min in 5% milk TBS-T, before 5x 5 min washing (∼10 mL in TBS-T) and incubation with secondary Rabbit-HRP antibodies (ab79773, Abcam) for 60 min. After more washes, blots were visualized using HRP after 2 min of incubation.

### Sortase labelling

Fluoresceine-amidite (FAM)-labeled peptide (FAM-HHHHHHLPETG) was added N-terminally to TXNL1 and the Eos-titin-tail substrate using Sortase ligation at room temperature for 30 min. Proteins (30-100 μM) were incubated in 50 mM HEPES, pH 7.4, 100 mM NaCl, 10 mM CaCl_2_, 0.5 mM TCEP with 500 μM peptide and 25 μM Sortase in 250-450 uL reaction volumes. Labelled proteins were bound to Ni-NTA, washed several times with 60 mM HEPES pH 7.4, 25 mM NaCl, 25 mM KCl, 5% glycerol, 10 mM MgCl_2_, 20 mM imidazole, and eluted with the same buffer supplemented with 250 mM imidazole. Labelled proteins were concentrated and fractionated by gel filtration using an SD75 increase 10/300 in 60 mM HEPES pH 7.4, 25 mM NaCl, 25 mM KCl, 5% glycerol, 10 mM MgCl_2_ to remove excess peptide. Labelled substrates were concentrated, snap frozen in liquid N_2_, and stored at -80°C as 10 μL single-use aliquots. For TXNL1 constructs, 0.5 mM TCEP was included throughout the purification and in the storage buffer.

### Insulin reduction assay

When reduced, the B chain of insulin will self-aggregate and precipitate, leading to increased turbidity as measured by absorbance at 650 nm. Insulin (I0516, Sigma Aldrich) at ∼1.7 mM was diluted to 30 μM in reaction buffer (60 mM HEPES, pH 7.4, 150 mM NaCl, 5% glycerol with 1 mM DTT, unless otherwise stated) with TXNL1 protein. A_650nm_ was measured every 30 seconds for at least 2 hours at 30°C.

### Substrate ubiquitination and degradation

2X reactions of His-mE1 (1 μM), Ubch7 (10 μM), Rsp5 (6 μM) and Ubiquitin (200 μM) were incubated with Eos-titin-tail substrate (5 μM) for ∼30 mins at room temperature in ubiquitination reaction buffer (60 mM HEPES, pH 7.4, 25 mM NaCl, 25 mM KCl, 10 mM MgCl, 0.5 mM TCEP, 5 mM ATP). Ubiquitinated Eos-titin-tail substrate was mixed with 2X concentrated hs26S proteasome (which was treated before with 2.5 μM Ub-prg for 30 mins) in a pre-warmed 384-well plate, and degradation was measured in a BMG Labtech CLARIOstar plate reader at 30°C by monitoring the emission at 520 nm after excitation with at 500 nm. To monitor formation of peptide products by SDS-PAGE analysis, N-terminally FAM-labelled ubiquitinated Eos-titin-tail substrate was used, and samples were boiled prior to gel loading to ensure Eos was fully unfolded and no longer fluorescence.

### ^FAM^TXNL1 proteasome binding assay

FAM-labeled TXNL1 and hs26S proteasome were incubated at 2X concentration, such that once diluted 1:1 with buffer or substrate, their final concentrations were at 50 nM for TXNL1 (unless otherwise stated) and 150 or 500 nM for hs26S proteasome. FAT10 substrate was prepared for degradation by incubating the FAT10 with Nub1 at 1:1.2 ratio for at least 15 min at 2 X concentration. Reactions were initiated by adding buffer or the pre-incubated substrate to TXNL1 with or without hs26S proteasome, and fluorescence polarization was recorded by measuring the emission at 535 nm after excitation at 480 nm in a preheated (30 °C) 384-well black plate (Costar) using a BMG Labtech CLARIOstar plate reader.

### Cryo-EM Sample preparation and data collection

Human 26S proteasome (4 μM, high-salt washed, see proteasome purification method) with recombinant FLAG-TXNL1 (16 μM) was diluted 1:1 with preincubated FAT10-Eos (12 μM) and NUB1 (16 μM) for ∼ 105 s before cryo-plunging. Sample buffer used was 20 mM HEPES pH 7.4, 25 mM NaCl, 25 mM KCl, 5 mM MgCl_2_, 2 mM ATP, 2.5% glycerol and 0.02% NP-40. Final samples had 2 μM 26S proteasome, and 3.5 μL were applied to glow discharged (glow discharging: 25 mA, 25 seconds) UltrAufoil^®^ R 2/2, 200 Mesh, Au grids (Q250AR2A, Electron Microscopy Sciences). Grids were plunge frozen in liquid ethane (cooled with liquid nitrogen) using a Vitrobot (ThermoFisher) at 12°C with 3 s of blot time. Data were collected as previously described ^25^, where briefly, clipped grids were transferred to a Titan Krios transmission electron microscope operated at 300 KeV (ThermoFisher) equipped with a Gatan K3 and an energy filter (GIF quantum). Images were taken using SerielEM at a nominal magnification of x81,000 (1.048 Å pixel size) in super resolution mode with a defocus ranging from −0.5 – 1.7 μm. We collected 50 frames per shot with a total electron dose ∼50 e^−^ Å^−2^s^−1^. A total of 9,239 movies were collected for the high-salt washed proteasomes with excess TXNL1 and 2 min after addition of the NUB1/FAT10-substrate. The dataset for low-salt washed proteasomes with sub-stoichiometric amounts of TXNL1 and at 30 s after substrate addition was previously collected ^25^.

### Cryo-EM data processing

Data processing was performed using CryoSparc v4.4 ^45^ and datasets were processed separately. Movies were corrected with patch motion and patch CTF, before blob picking and particle extraction with a box size of 600, but binned by 2x. Particles were sorted by multiple rounds of 2D classification before *ab initio* 3D reconstruction and heterogenous refinement with 10 classes and a refinement box size set to 128. Each class was refined by homogenous refinement and high-resolution classes of 26S and 30S proteasomes were selected, pooled, and extracted with a 600-box size and no binning. All particles were refined using non-uniform refinement in C2.Particles were symmetry expanded in C2 followed by recentering, such that the proteasome regulatory particle is in the center of the box, and subsequently extracted with a box size of 340. Particles were then subjected to 3D-reconstruction to ensure their correct position and subjected to heterogenous refinement with 10 classes. Classes were pooled based on the global conformation of the RP, either belonging to resting state (RS) or processing states (PS) and refined using non-uniform refinement.

To separate processing-state proteasomes into individual states, a generous mask for the RP was generated for local refinement and aligned particles were subject to alignment-free 3D classification (10 classes, filtered at 6 Å), from which 4 dominant classes emerged. A mask for the ATPase domains was then generated for each class, followed by local refinement and another round of 3D classification (10 classes, filtered to 6 Å). Each of the 10 classes was refined with local refinement and similar states (no obvious differences) were combined, such that 6 ATPase motor conformations were achieved. ATPase 3D classification for PS_Rpt5_ resulted in different classes with varying positions of the seam subunit Rpt4 and the PS _Rpt1_. For PS_Rpt6_ multiple poor-quality particles were included likely due to lower resolution. Therefore, another round of heterogenous refinement and local refinement of the ATPase was performed and resulted in a relatively high resolution PS_Rpt6_ conformation, while the rest of the particles appears to be junk or low-resolution data. For each processing conformation an RP mask was generated, and local refinement was used to generate final unsharpened maps. An ATPase motor mask was used for processing maps from the 30 second dataset ^25^. While we describe distinct states of the 26S proteasome, it is important to note that alignments of particles and 3D reconstructions can be dominated by more rigid parts of a protein complex. In addition,within each discretestate, there is continuous motion present throughout the 26S proteasome complex, and this can vary depending on the defined conformational state described. For example, Rpn10 and Rpn1 can be at lower resolutions relative the AAA+ motor. It should be noted that due to current classification techniques particles are sorted based on their probability of belonging to each assigned state, and more discrete states might be observed when increasing particle numbers. Although several groups previously generated composite maps from individually locally refined maps ^18,20^, we did not use this approach. However, for guidance in model building and map representation and to aid with the anisotropy in map resolution, we used maps sharpened with deepEMhancer ^46^, but always validated model interpretations with unprocessed maps and locally refined maps.

For the 30-second dataset with partial occupancy of TXNL1 on Rpn11, we also generated masks that encompassed TXNL1, Rpn2, Rpn11 and Rpn10 and used local refinement to improve resolution. Using these locally refined maps, alignment-free 3D classification was used followed by local refinement of each separate class based on the presence of absence of TXNL1. Particles were pooled into two groups, with and without TXNL1, and subjected to local refinement using the same mask for TXNL1. The particle numbers for the presence and absence of TXNL1 were used to roughly calculate the ratio of TXNL1 binding to a particular processing-state conformation. However, it should be noted that numbers are estimates due to lower resolutions in certain classes and particles being sorted based on the probability of belonging to a certain class.

For the resting-state proteasome, RP-aligned particles were subjected to alignment-free 3D classification, from which 4 major classes emerged. Each class was refined by non-uniform refinement, whereby Class1 was considered junk due to broken RP, Class2 was characterized by a flexible Rpn1, and Class3 and Class4 were charactered as high resting state conformations with a shift in the position of the RP when compared. Class 2 was further processed by aligned local refinement with a mask encompassing Rpn10, Rpn2, Rpn11 and the density in between. Aligned particles were subject to alignment-free 3D classification, which generated 10 classes of varying resolutions. We chose 3 classes with interesting features to further refine. Two classes showed the PITH domain of TXNL1 in forward and backward conformations and were refined through local refinement with a whole RP mask. One conformation appeared similar to so-called ‘deubiquitination states’ E_A2_ and E_B_ ^19^ and showed density near Rpn11, which is likely a ubiquitin-like domain. After local refinement with a whole RP mask, we low-pass filtered the whole RP-refined map to 6 Å and docked the atomic model for the E_B_ state (PDB ID 6MSE), using ChimeraX ^47^. Local refinement with a mask encompassing Rpn11, the extra density, the coiled coil of Rpt4/5, Rpn10 and Rpn2, and subsequent 3D classification did not further improve the resolution. For Class 3 and Class4 with a slight shift in the RP, there was an ensemble of density where we would expect the PITH domain of TXNL1. We kept Class3 and Class4 separate and used local refinement to align particles using a mask encompassing Rpn2, Rpn10, Rpn11 and the ensemble of density in between those subunits. CryoSparc 3D variability and cluster analysis ^48^ was used to separate out particles based on the position of the PITH domain. We could characterize two major conformations, a forward and backward conformation. Those states, named RS.1 TXNL1 state 1, RS.1 TXNL1 state 2, RS.2 TXNL1 state 1, and RS.2 TXNL1 state 2, were refined by local refinement with a whole RP mask, and maps sharpened with deepEMhancer ^46^ along with unprocessed maps were used for representing density.

### Cryo-EM model building and model visualization

Previously built atomic models for human 26S proteasomes ^25^ were used as starting models for each proteasome structure. Atomic models were generated for resting-state proteasome maps: RS.1 TXNL1 state 1, RS.1 TXNL1 state 2, RS.2 TXNL1 state 1, and RS.2 TXNL1 state 2. Briefly, models were fit using ChimeraX Fit to map and individual subunits were manually adjusted using the same function. TXNL1’s PITH domain was truncated from a model generated by AlphaFold ^49^. Phenix refine ^50^ with simulated annealing and morphing turned on for the first round was used to adjust models, followed by several rounds of iterative model building in Coot ^51^ and real space refinement in Phenix without simulated annealing and morphing. For atomic models of the processing-state proteasomes, deepEMhancer maps were used to guide model building. For AAA+ domains, Rpts were separated into multiple parts and rigid bodies using ChimeraX Fit to map, followed by several rounds of manual adjustment in Coot. Once roughly assembled and fit, models were subjected to multiple rounds of manual adjustment in Coot and real space refinement with Phenix (refinement was typically done using unprocessed maps). The chromophore for Eos (code CR8 in Coot) was fit and adjusted in Coot, followed by Phenix real space refinement using a restraints cif file generated in Phenix. PyMol (The Molecular Graphics 981 System, Version 1.8, Schrödinger, LLC; http://www.pymol.org/), UCSF chimera ^52^ and ChimeraX ^47^ were used to generate figures. Local resolutions were calculated using CryoSparc local resolution estimation with 0.143 from FSC curves and displayed on locally filtered maps (using a B-factor obtained during refinement) and displayed using ChimeraX with color surface.

## RESOURCE AVAILABILITY

## Lead contact

Further information and requests for resources and reagents should be directed to and will be fulfilled by the lead contact, Andreas Martin.

## Materials availability

All constructs generated in this study are available from the lead contact upon request and completion of a Material Transfer Agreement.

## Data and Code Availability

- All data generated or analyzed during this study are included in this manuscript and the Supplementary materials. The cryo-EM density maps and corresponding atomic coordinates for human 26S proteasomes actively degrading FAT10 bound to TXNL1 can be found on the Electron Microscopy Data Bank (EMDB) under the following EMDB accession codes and under Protein Data Bank (PDB) accession codes: EMD-47719/PDB-9E8G (PS_Rpt5_), EMD-47724/PDB-9E8L (PS_Rpt4_), EMD-47725 /PDB-9E8N (PS_Rpt3_), EMD-47723/PDB-9E8K (PS_Rpt6_), EMD-47726/PDB-9E8O (PS_Rpt2+Eos_), EMD-47722/PDB-9E8J (PS_Rpt1_), EMD-47721/PDB-9E8I (RS.1_TXNL1 Forward_) and EMD-47720/PDB-9E8H (RS.2_TXNL1 Backward_), EMD-47727/PDB-9E8Q (PS_Rpt2_).
- This paper does not report original code.
- Any additional information required to reanalyze the data reported in this paper is available from the lead contact upon request.

## Acknowledgments

We thank all members of the Martin lab for discussion and support. We thank Lan Huang (UC Irvine) for gifting httb-HEK293 cells and the UCB Cell Culture Facility (RRID: SCR_017924) for maintaining the httb-HEK293 cell culture. Cryo-EM data were collected at the UCB Cal-Cryo facility, and we thank Dan Toso and Ravindra Thakkar for cryo-EM operational support.

## Author contributions

C. A. and A.M. conceived the study, C.A. and A.M. designed experiments, C.A. cloned constructs, expressed, and purified proteins, and C.A., Z.Z. and K.C.D. performed biochemical measurements as well as data analyses. C.A. performed cryo-EM sample preparation, data collection, and data processing. C.L.G. and C.A. generated atomic models based on cryo-EM maps. C.A. and A.M. wrote the manuscript with comments from all authors.

## Extended Figures

**Extended Figure 1:**
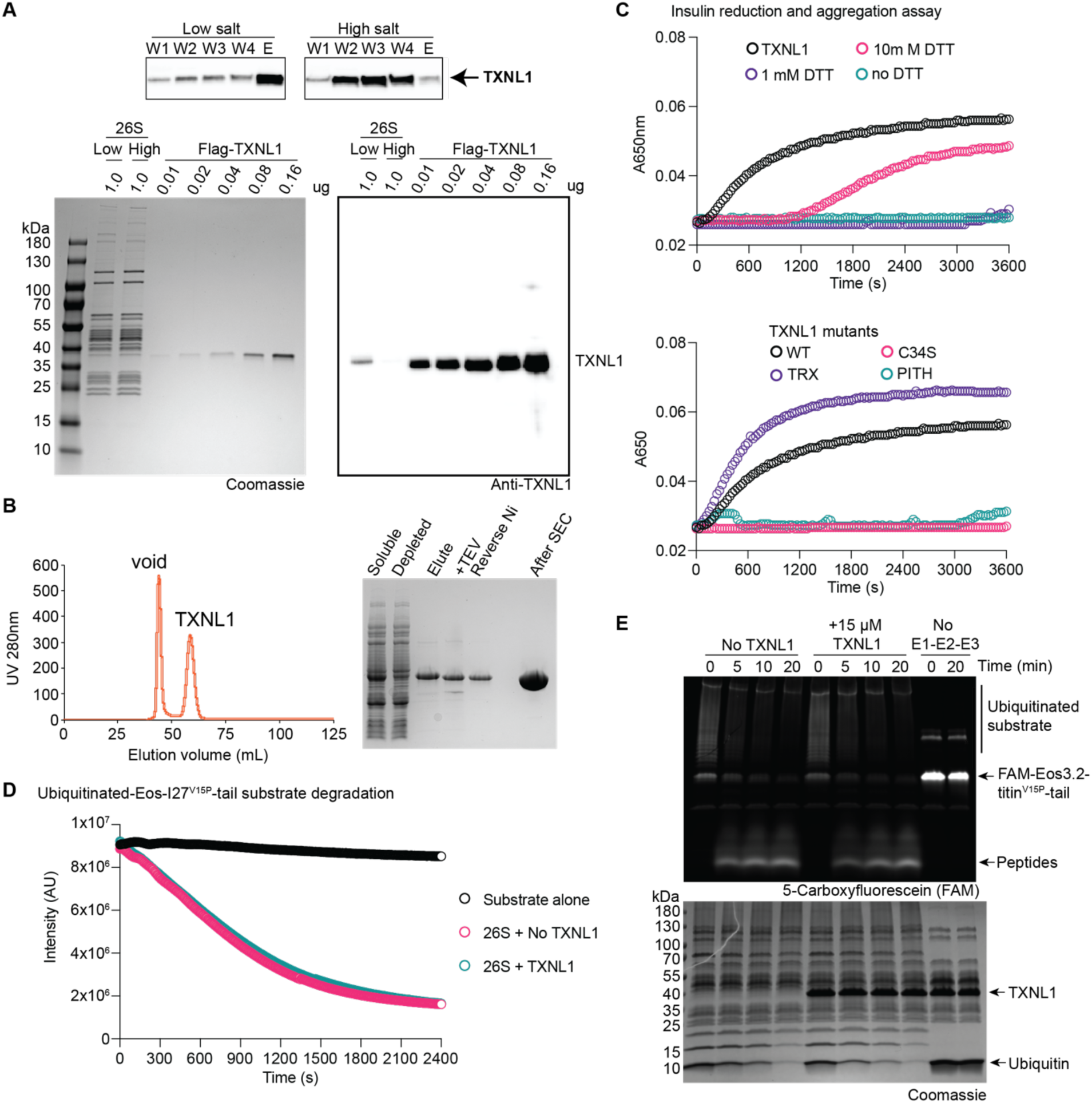
TXNL1 is redox active and co-purifies with human 26S proteasomes. A) Purification of TXNL1-bound and TXNL1-free human 26S proteasomes from HEK293 cells. Top: Western blots showing aliquots from the washing (W1-W4) and elution (E) steps in low-salt or high-salt buffer for HTBH-tagged proteasomes that were immobilized on streptavidin agarose. Bottom left: Coomassie-stained SDS-PAGE gel showing the separation of 1 μg human 26S proteasomes purified by size-exclusion chromatography after previous low salt or high salt washes and compared to specific concentrations of recombinant FLAG-tagged TXNL1 purified from *E. coli.* Bottom right: Western blot of the SDS-PAGE samples on the left, showing TXNL1 levels that co-purified with low-salt or high-salt washed proteasomes in comparison to recombinant His-FLAG-tagged TXNL1. Low-salt washed proteasomes contain sub-stoichiometric amounts of TXNL1, whereas TXNL1 levels for high-salt washed proteasomes are almost undetectable. B) Left: Elution profile for the size-exclusion chromatography (SD75 16/600) of recombinant human TXNL1 that was expressed in *E. coli* and Ni-NTA affinity purified using its His-(TEV)-FLAG tag. Right: Coomassie-stained SDS-PAGE gel with aliquots from individual stages of recombinant TXNL1 purification. C) Redox activity of recombinant TXNL1 (15 μM) measured by the increase in turbidity (absorbance at 600 nm) that results from the reduction and consequent aggregation of insulin (30 μM). Activities are compared to different concentrations of DTT (top) and TXNL1 mutants that contained only the N-terminal catalytic TRX domain, the C-terminal PITH domain, or the C34S mutation in the catalytic CXXC motif (bottom). D) Degradation of Eos-titin^V15P^-tail (5 μM) substrate by human 26S proteasome (200 nM) in the absence or presence of excess TXNL1 (15 μM), monitored by the loss of Eos fluorescence. E) FAM fluorescence detection (top) and Coomassie staining (bottom) of the SDS-PAGE gel with samples from the degradation of N-terminally FAM-labeled and ubiquitinated ^FAM^Eos-titin^V15P^-tail substrate (2.5 μM) by human 26S proteasomes (200 nM) in the absence and presence of excess TXNL1 (15 μM). The right 2 lanes show a negative control with non-ubiquitinated substrate (no E1, E2, E3 enzymes).

**Extended Figure 2:**
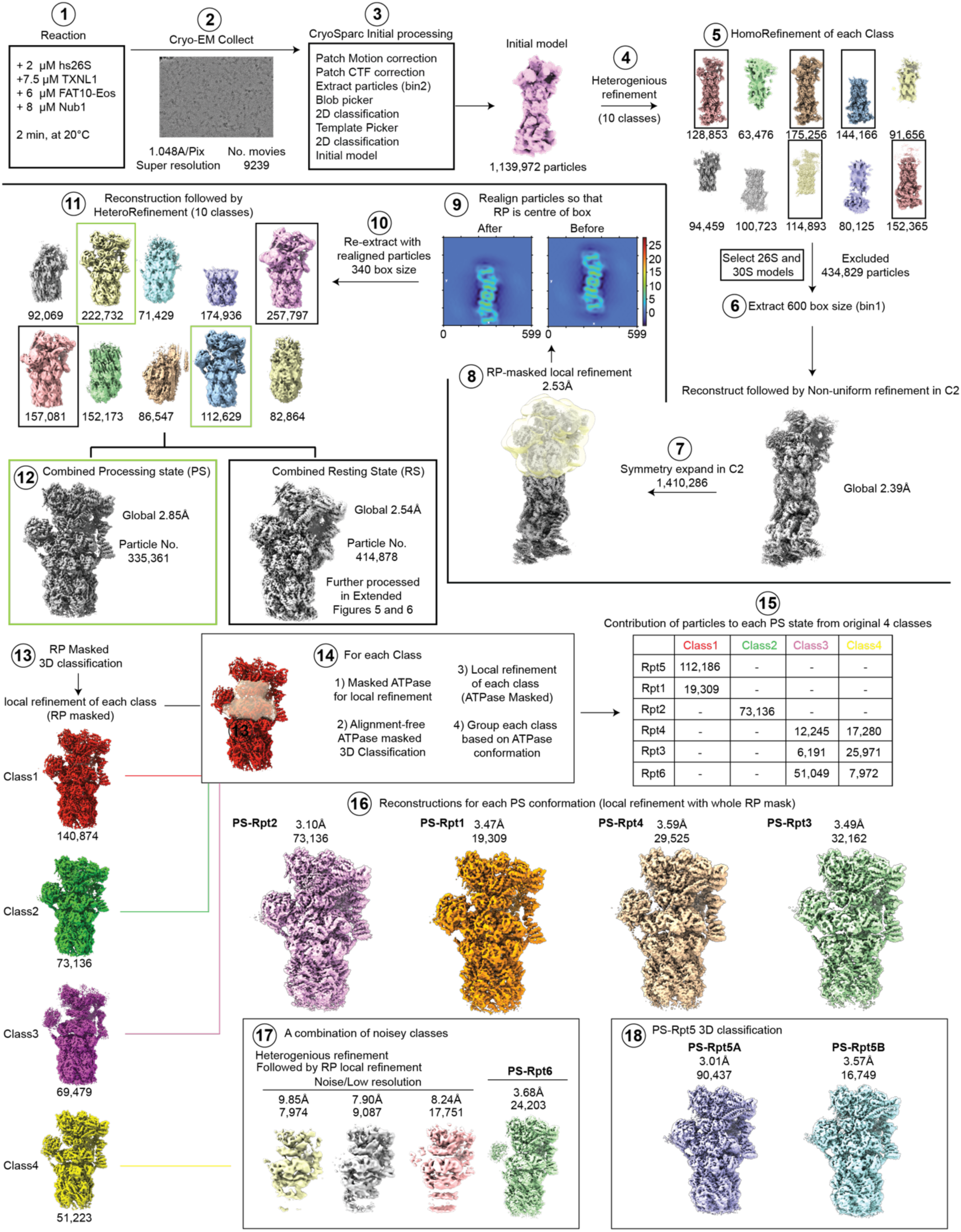
Workflow for the cryoEM structure determination of human 26S proteasomes. Shown are the sequential steps taken to process cryo-EM data from movies to final maps for the human 26S proteasome at ∼ 2 min after incubation with excess Txnl1, FAT10-Eos substrate, and the NUB1 cofactor. From step 13 the workflow focuses on proteasomes in the processing states (PS), while the subsequent workflow for proteasomes in the resting state (RS) is shown in Extended Figures 5 and 6.

**Extended Figure 3:**
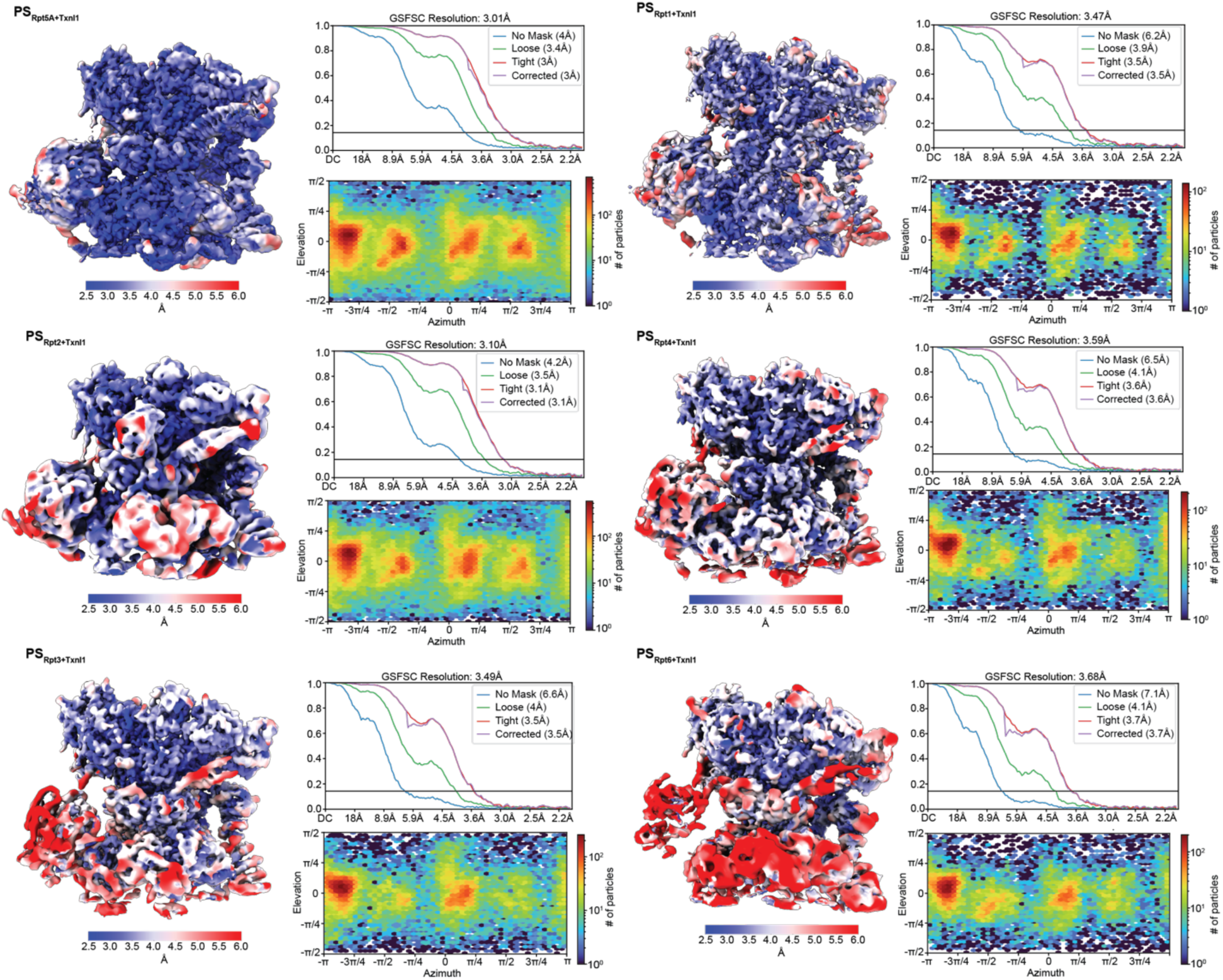
Local resolution maps for the 6 substrate-processing (PS) conformations of the human 26S proteasome in the presence of excess TXNL1. These maps resulted from the workflow shown in Extended Figure 2. PS states are named based on the Rpt subunit that resides at the top of the substrate-engaged spiral staircase. Local resolutions were calculated with CryoSparc using a FSC cutoff of 0.143.

**Extended Figure 4:**
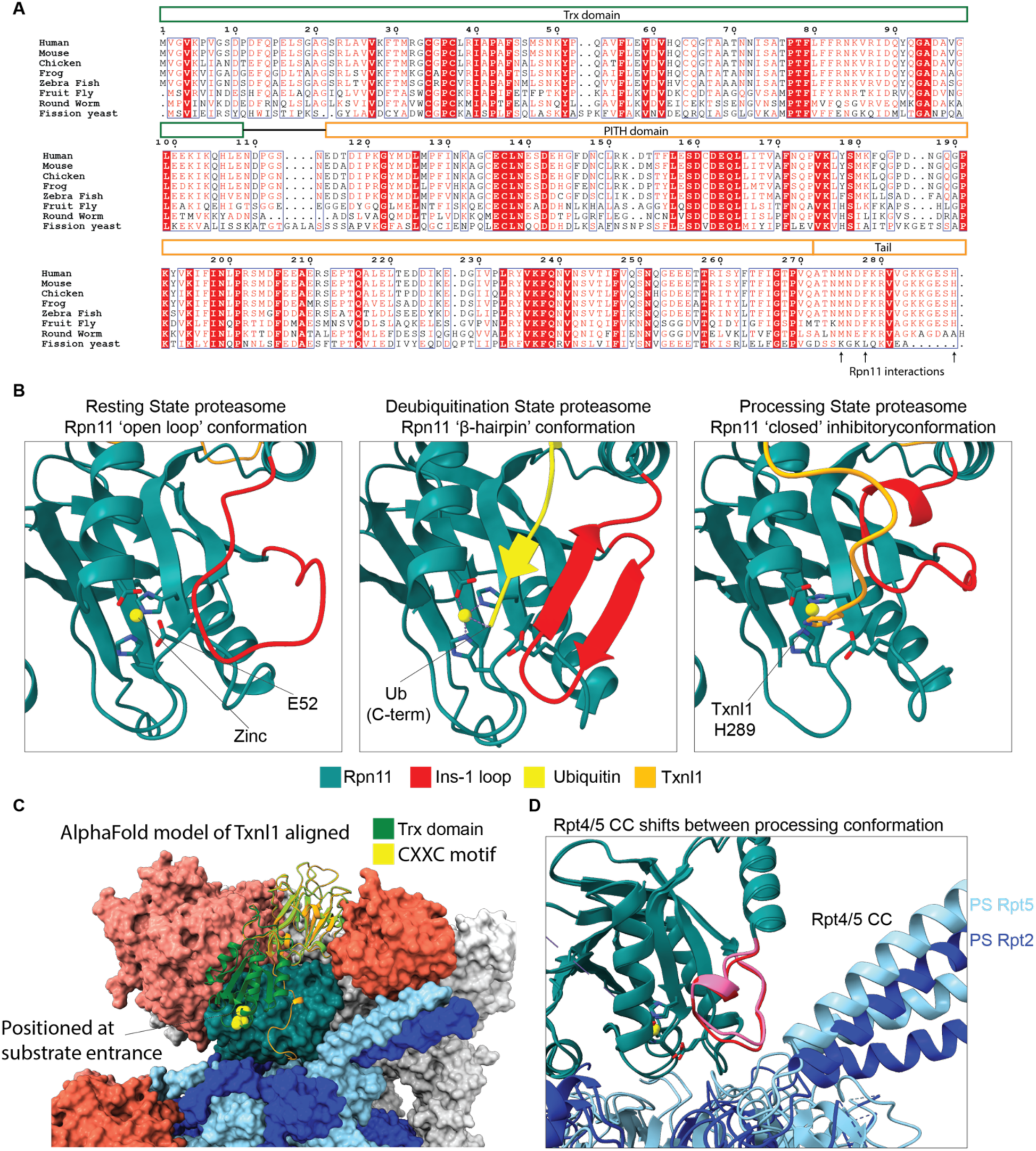
TXNL1 interacts with Rpn11 in a conformation-specific manner using its conserved C-terminal tail. A) B) Sequence alignment of TXNL1 from uniport sequences using ENDscript at default settings: *Homo sapiens* (Q43396), *Mus musculus* (Q8CDN6), *Gallus gallus* (A0A1D5NUH1), *Xenopus tropicalis* (F6RTD9), *Danio rerio* (F6NTA0), *Drosophila melanogaster* (Q9VRP3), *Caenorhabditis elegans* (G5EES9) and *Schizosaccharomyces pombe* (Q9USR1). Red underlay shows conserved residues and blue boxes indicate similarity in amino acid identity. Residues in the C-terminal tail that mediate the interaction with Rpn11, including the terminal His, are highly conserved, except for fission yeast. B) Atomic models of Rpn11 with the Insert-1 (Ins-1) region in three conformations. The open loop conformation is found in resting-state proteasomes, the deubiquitination conformation with a β-hairpin is adopted upon ubiquitin binding to Rpn11, and the inhibitory, closed conformation is observed substrate-processing proteasomes. The three models are derived from our structure for the resting state RS.1 with TXNL1 in the forward orientation, the crystal structure of ubiquitin-bound Rpn11-Rpn8 (PDB ID: 5U4P), and our structure of the substrate-processing state PS_Rpt5_, respectively. C) Space filling atomic model of PS_Rpt5_ with the ribbon-represented PITH domain of TXNL1 in orange. The AlphaFold model for full-length TXNL1 is aligned by its PITH domain (light green) with the PITH domain in our structure and shows the TRX domain (dark green) with its catalytic CXXC motif (yellow spheres) close to the entrance of the ATPase motor. D) Overlay of the atomic models for PS_Rpt5_ (Rpts in cyan) and PS_Rpt2_ (Rpts in dark blue), aligned by Rpn11 (petrol), shows a movement of the Rpt4/Rpt5 coiled coil that significantly increases the gap to the Ins-1 loop (pink for PS_Rpt5,_ red for PS_Rpt2_).

**Extended Figure 5:**
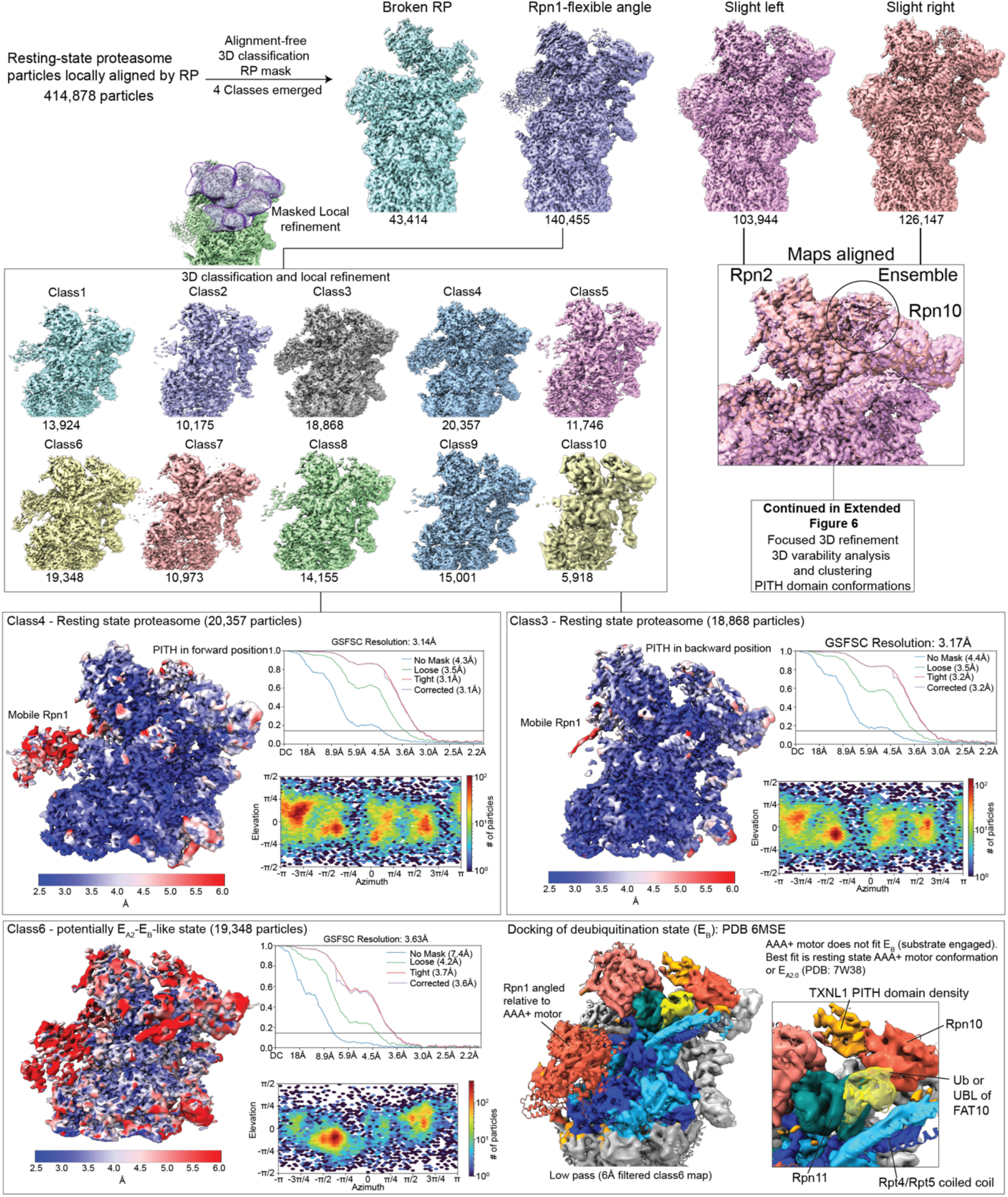
Further cryo-EM data processing of resting-state (RS) proteasomes in continuation from extended Figure 2. Classification leads to the emergence of 4 classes. Two of these classes, called ‘Slight left’ and ‘Slight right’ based on the position of the regulatory particle, show an ensemble of PITH domain densities and were further processed as described in Extended Figure 6. Further classification of class 2 with variability in the Rpn1 subunit revealed 10 additional classes at moderate resolution. Three of them, class 4, class 3, and class 6 are shown in more detail below. Class 4 has TXNL1’s PITH domain in the forward conformation, whereas class 3 shows the PITH domain oriented backwards. Class 6 represents a unique class with extra density on Rpn11 that may originate from ubiquitin or a ubiquitin-like domain, for instance of FAT10. A human proteasome structure (PDB: 6MSE) with ubiquitin-bound Rpn11 was used for comparison, to color the class 6 map, and show similarities.

**Extended Figure 6:**
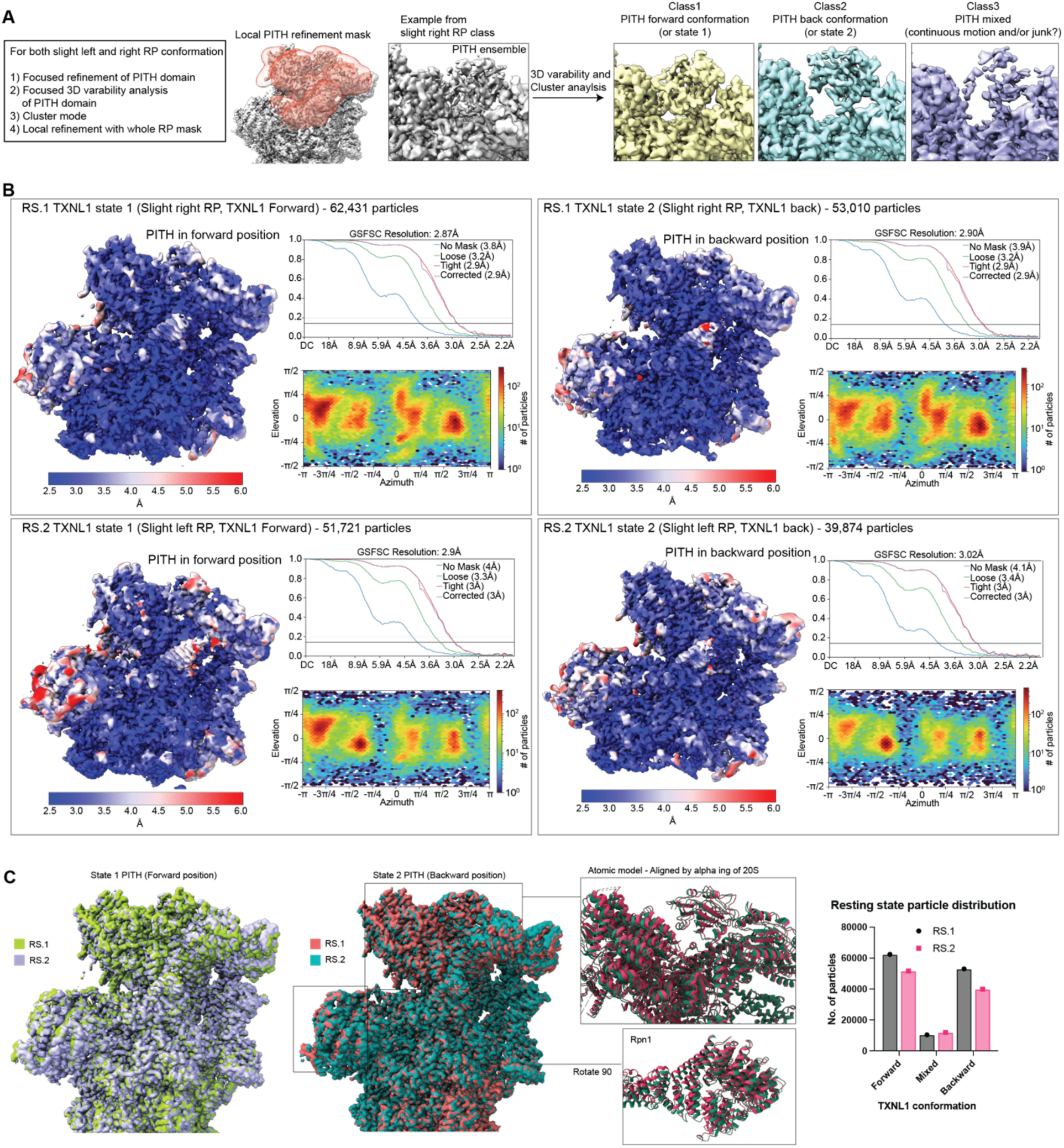
Further cryo-EM data processing of resting-state (RS) proteasomes with bound TXNL1 in continuation from extended Figure 5. A) ‘Slight left’ and ‘Slight right’ particles were processed separately. Particle stacks were aligned (local refinement in CryoSparc) with a local mask surrounding the PITH domain, Rpn2, Rpn10, Rpn11, and the N-ring of the Rpt hexamer. CryoSparc 3D variability analysis of aligned particles with the local mask, followed by 3D cluster analysis, allowed grouping of particles into two distinct conformations, forward and backward with respect to the PITH domain position. A third group of particles showed the PITH domain in variable positions and could not be assigned to a particular state, suggesting that the PITH domain has continuous motion between forward and backward orientations. B) Local refinement with a mask surrounding the whole RP resulted in 4 distinct conformations defined by a slight shift in the RP and the two orientations of the PITH domain: RS.1 TXNL1 Forward (RS.1 state 1), RS.1 TXNL1 Backward (RS.1 state 2), RS.2 TXNL1 Forward (RS.2 state 1), and RS.2 TXNL1 Backward (RS.2 state 2). Each structure was resolved to high resolution, as demonstrated by the maps colored according to estimations of local resolutions, calculated with CryoSparc using a 0.143 FSC cutoff. C) Alignments of the atomic models for RS.1 and RS.2, states 1 and 2, highlight the shift in the RP, in addition to variations in Rpn1’s position. Right: Particle numbers for each conformational state that were used to calculate the percentages of proteasomes with TXNL1 in the forward, backward, or mixed orientations shown in Figure 2B.

**Extended Figure 7:**
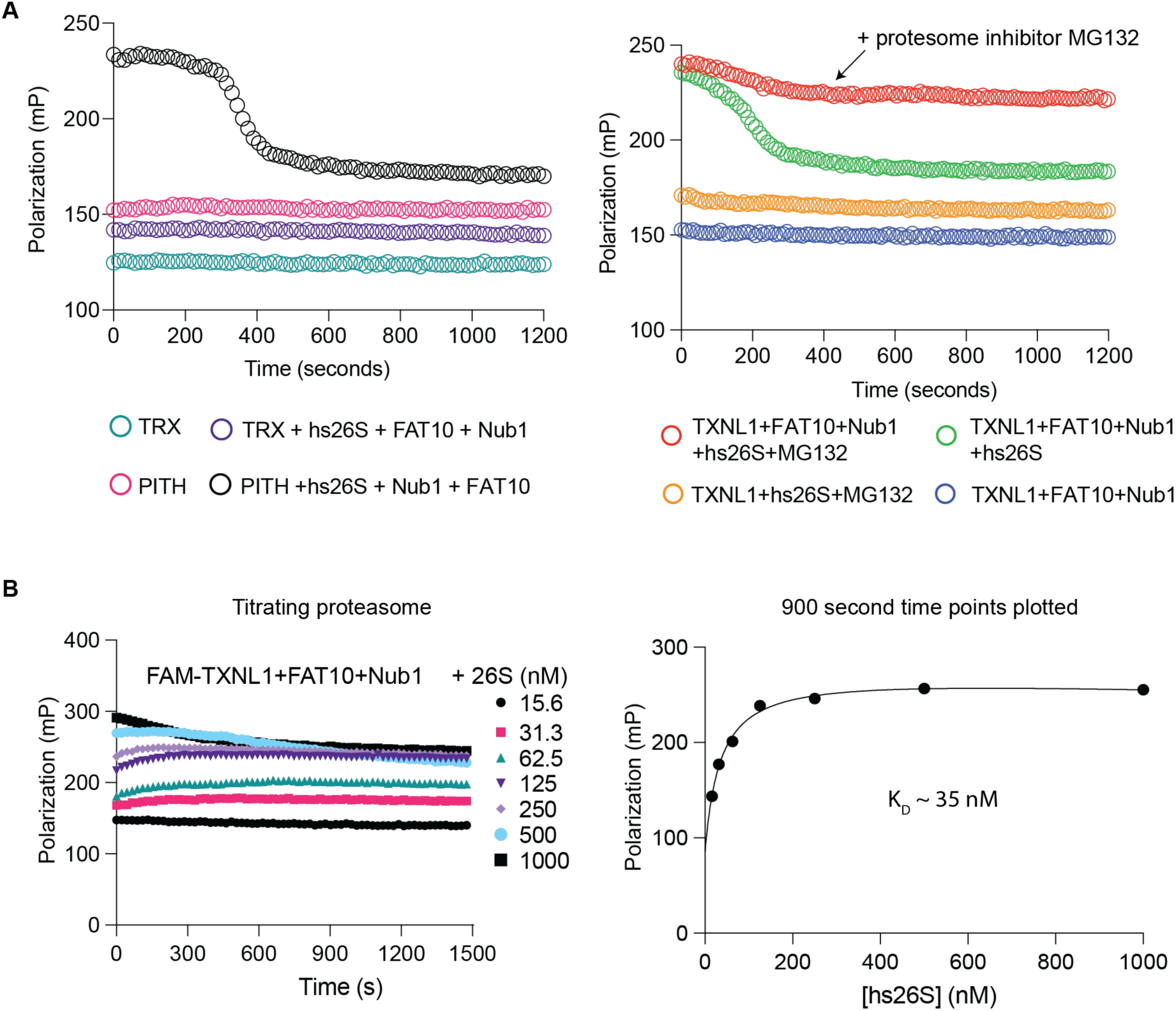
TXNL1 dissociates from proteasomes as they transition back from processing to the resting state, but remains bound when degradation is stalled. A) Left: TXNL1’s PITH domain is necessary and sufficient for the conformational selective binding to actively degrading 26S proteasomes. Shown are representative traces for the fluorescence polarization of FAM-labeled TRX domain (50 nM) or PITH domain (50 nM) that was incubated in isolation or with human 26S proteasome (500 nM) in the presence of 10 μM FAT10 substrate and 12 μM NUB1 cofactor. Right: Proteasomes whose proteolytic cleavage of the FAT10 substrate was inhibited by MG132 and which were therefore stalled during substrate translocation interact stably with TXNL1. Shown are the fluorescence polarization time courses for N-terminally FAM-labeled full-length ^FAM^TXNL1 (50 nM) that was incubated with FAT10 (5 μM), NUB1 (6 μM), and human 26S proteasome (500 nM) in the absence or presence of MG132 (20 μM), as indicated. B) Titration of proteasomes that were stalled during substrate degradation through MG132 inhibition of proteolysis reveals the TXNL1 binding affinity for processing-state proteasomes. Left: Shown are the fluorescence polarization traces for ^FAM^TXNL1 (50 nM) incubated with human 26S proteasome at indicated concentrations in the presence of FAT10 substrate (5 μM) and NUB1 cofactor (6 μM). Right: ^FAM^TXNL1 binding curve derived from the 900 s time points of the polarization traces shown on the left indicates an affinity of *K_D_* ∼ 35 nM.

**Extended Figure 8:**
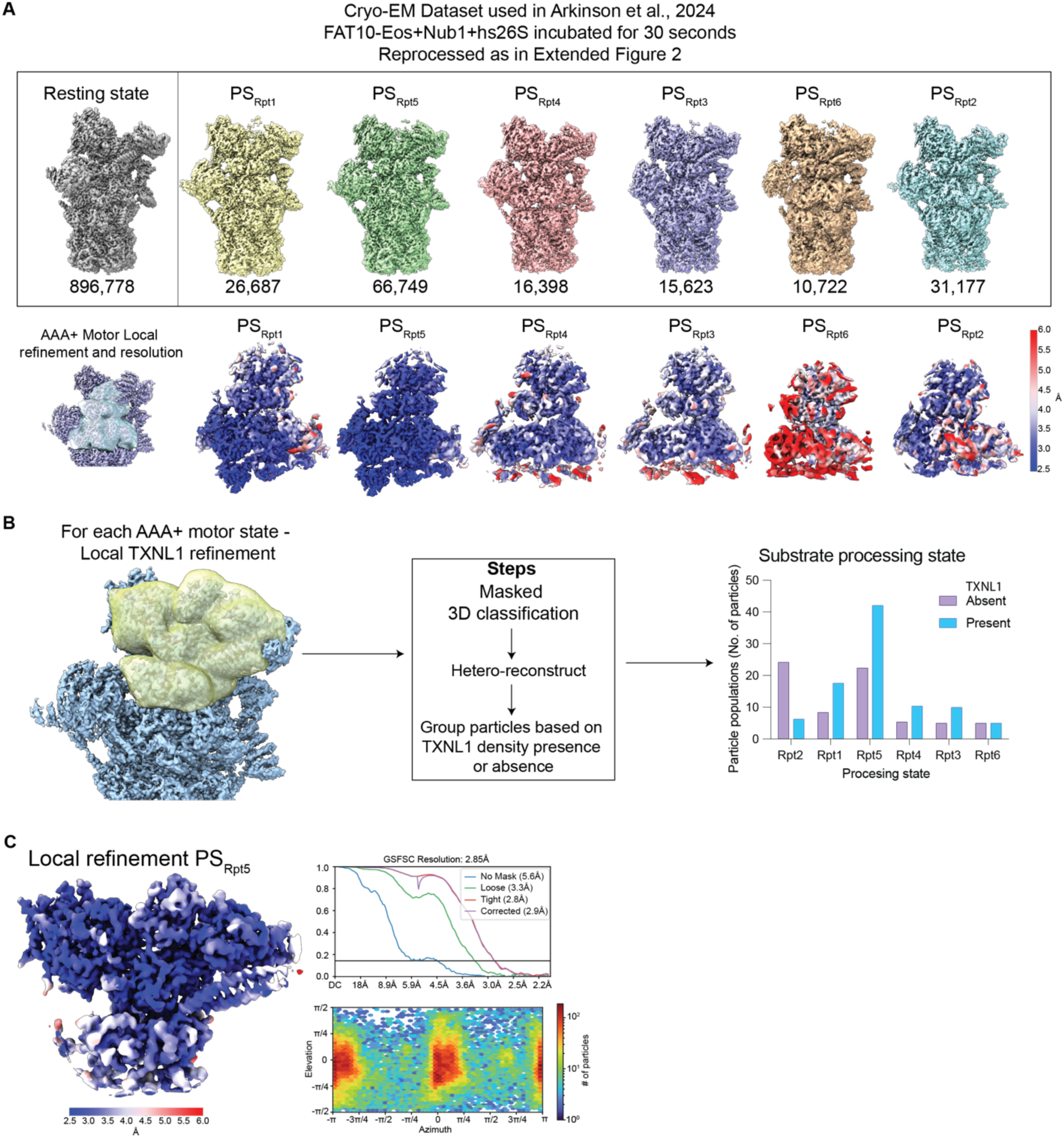
Cryo-EM data processing for the human 26S proteasomes 30 s after incubation with FAT10 substrate and NUB1 cofactor. A) Data were processed as shown in the workflow in Extended Figure 2. Top: Shown are the unsharpened maps and particle numbers for each state. Bottom: To improve resolution of the AAA+ ATPase domains a local mask was generated for the Rpt hexamer and Rpn11 and local refinement was used. Maps show local resolutions of the Rpt subunits and Rpn11 for each processing state. B) Human proteasomes used in this dataset were purified from HEK293 cells under low-salt conditions and therefore contain sub-stoichiometric amounts of bound TXNL1. Extensive local 3D classification focused on TXNL1 for each processing state allowed separation of particles with and without bound TXNL1. Briefly particles were locally aligned by a mask surrounding TXNL1’s PITH domain and subjected to 3D classification with a filtered resolution of 10 Å. Each class was reconstructed and refined to group particles based on the presence or absence of PITH density, and the particles numbers for PITH-bound and -unbound proteasomes are plotted for each processing state (right). C) Local resolution of locally refined PS_Rpt5_ after removing unbound TXNL1 particles.

**Extended Figure 9:**
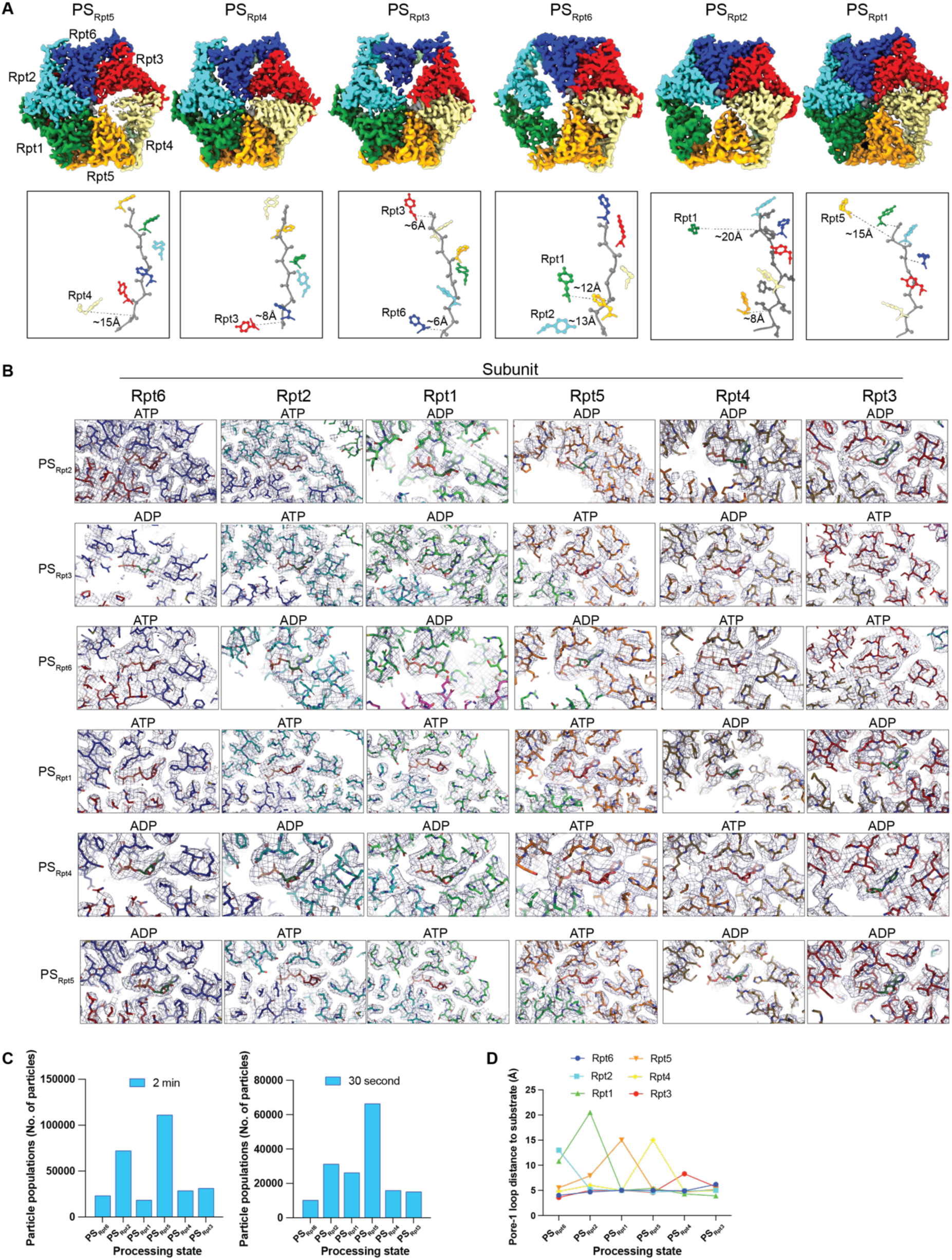
Comparisons of the six discrete substrate-processing states for the human 26S proteasome regarding spiral-staircase arrangements, pore-loop interactions with the substrate polypeptide, nucleotide occupancies, and asymmetric particle distributions. A) Top: Cryo-EM densities of the AAA+ motor for all processing-state (PS) conformation, colored by Rpt subunit. Substrate-disengaged seam subunits are at lower resolution likely due to higher mobility and various vertical positions between the bottom and top of the spiral staircases. PS_Rpt3_, PS_Rpt6,_ and PS_Rpt2_ show larger density gaps between subunits at the seam, likely because they contain two substrate-disengaged seam subunits with potentially high continuous motions. Bottom: Spiral staircase arrangements of pore-1-loop Tyr residues (Phe for Rpt5) for each processing state. Distances between the substrate backbone and the backbone Cα of the pore-1 loop aromatic residue are indicated for the disengaged seam subunits. Interestingly, some seam subunits reside at the bottom, while others are observed toward the top of the staircase. Positions of the pore-loop residues for these seam subunits are approximate due to the higher mobility and consequently lower resolution. B) Cryo-EM densities and atomic models, including the specific nucleotide (ATP or ADP) for all Rpts of the six processing states. C) Particle numbers observed for each processing state in the separate cryo-EM data sets for the 26S proteasome in the presence of sub-stoichiometric TXNL1, 30 s after incubations with the FAT10 substrate (left) or in the presence of excess TXNL1, 2 min after incubation with FAT10 (right). D) Plotted distances between the substrate backbone and the backbone backbone pore loop interactions with substrate backbone and the backbone Cα of the pore-1 loop aromatic residue for Rpt1-6 in each substrate-processing state of the proteasome.

